# Multiple scales of coordination along the body axis during *Drosophila* larval locomotion

**DOI:** 10.1101/2025.08.21.671596

**Authors:** Marie R Greaney, Ellie S Heckscher, Matthew T Kaufman

**Affiliations:** Committee on Neurobiology, The University of Chicago, Chicago, IL, United States; Department of Molecular Genetics and Cell Biology, The University of Chicago, Chicago, IL, United States; Neuroscience Institute, The University of Chicago, Chicago, IL, United States; Department of Organismal Biology and Anatomy, The University of Chicago, Chicago, IL, United States; NSF-Simons National Institute for Theory and Mathematics in Biology

## Abstract

Coordinated movement along the body axis is critical to locomotion. In segmented, limbless animals, anterior (head) and posterior (tail) segments play different roles in locomotion, leading to a need for flexible coordination across body regions. Larval *Drosophila melanogaster* present a tractable experimental model for limbless, segmented crawling given the extensive genetic tools available and the optical clarity of the body. Prior work has suggested that, during crawling, all larval body segments contract similarly, despite the fact that each crawl cycle comprises two overlapping phases: a piston involving the most posterior segments, and a peristaltic wave involving all body segments. To test whether coordination varies regionally during locomotion, we expressed GCaMP in body wall muscles of larvae of either sex, and recorded segmental contraction kinematics and muscle recruitment during many cycles of locomotion in linear channels. Facilitated by machine vision techniques, we discovered new features of larval crawling at multiple scales. First, the propagation of both contraction and recruitment waves slowed approaching mid-body segments, then sped up towards the head. Second, the timing relationship between contraction and recruitment waves could be highly variable in anterior segments. Third, contraction durations showed particularly strong intersegmental correlations among posterior segments. These data suggest posterior segments coordinate to power the piston phase while anterior segments tolerate greater flexibility to enable reorienting behaviors. Our results depict an unanticipated degree of axial heterogeneity in the coordination of limbless crawling, opening new avenues to study the origins of whole-body coordination and the consequences of segmental diversity for locomotion.

**SIGNIFICANCE STATEMENT:** Most animals move through the world by coordinating muscle contractions across the body’s many segments. Studying how segments are coordinated in limbless animals can uncover principles that are fundamental for movement across the animal kingdom. However, in one of the best genetic model organisms to address this question, the larval fruit fly, it was unknown how the nearly identical body segments are coordinated across the body. We used machine vision techniques to discover an unanticipated degree of heterogeneity: the timings of muscle activity and movements are more coordinated among tail-end than head-end segments. This provides new insights into animal movements, and opens opportunities to study how interactions of the body and nervous system coordinate segmental movements across an entire animal.

## INTRODUCTION

The coordination of various movement features across body regions—timings, amplitudes, rates, and durations of contraction—is critical for effective locomotion. Coordination is well-characterized for certain types of locomotion—anguilliform swimming, vermiform crawling, and inchworm crawling—in certain limbless animals, including leech, earthworm, tadpole, and lamprey, whose bodies are composed of near-identical segments arrayed along the body axis. A general picture emerges that coordination of segmental movements can be highly consistent or can vary cycle-to-cycle, depending on environmental substrate (Gray et al., 1938; Kristan et al., 1974; Kahn et al., 1982; Wallén & Williams, 1984; Stern-Tomlinson et al., 1986).

Larval insects are becoming an important model for studies of behavior, neurophysics, motor control, development, and evolution. Insect larvae possess limbless, segmented bodies. They navigate through their various environments using a behavioral repertoire composed of alternating lateral head bends or turns and linear forward “visceral pistoning” strides (Simon et al., 2010; Green et al., 1983). However, the coordination of segmental movements during visceral pistoning remains poorly characterized.

Visceral pistoning locomotion comprises overlapping piston and wave phases that together constitute a stride cycle (Simon et al., 2010; Heckscher et al., 2012). During the piston phase, the animal’s tail end, head end and internal viscera move with near synchrony, translocating the animal’s center of mass, while the rest of the external body remains largely stationary. During the wave phase, each segment of the body wall contracts sequentially in a posterior-to-anterior peristaltic wave, returning the animal’s exterior into register with the interior.

Visceral pistoning has been modeled as uniform propagation of a contraction wave through a series of identical and identically coupled units (Gjorgjieva et al., 2013; Pehlevan et al., 2016; Loveless et al., 2019). Kinematic studies that sample a small number of cycles have suggested that body segments’ contractions are similar and have consistent timing relationships (Gjorgjieva et al., 2013; Pulver et al., 2015; Sun et al., 2022). However, evidence from dissected preparations reveals substantial differences in the intrinsic properties of segmental circuits along the body axis, particularly between posterior and anterior segments (Jonaitis et al., 2022). These intrinsic differences raise questions about the presumed identical nature of segments’ motor outputs, and the strength and spatial scale of segmental coordination.

The field has made major progress in understanding the neural control of visceral pistoning, but despite this, the behavior and how it relates to neuromuscular activity remain incompletely understood. Researchers have delineated neural components of the circuits that subserve visceral pistoning (e.g., Kohsaka et al., 2014; Berni, 2015; Fushiki et al., 2016; Kohsaka et al., 2019) and recorded nervous system activity in dissected preparations (Pulver et al., 2015; Tastekin et al., 2018; Jonaitis et al., 2022). However, the relationships between centrally-driven muscle recruitment and the kinematics of movement are not understood.

This study aimed to determine how segmental contraction and muscle recruitment are coordinated across the *Drosophila* larval body axis during visceral pistoning locomotion. To do so, aided by machine vision techniques, we recorded >3000 cycles of forward visceral pistoning from larvae expressing calcium indicators in all body wall muscles as they crawled inside linear channels. In visceral pistoning locomotion, unlike other forms of limbless locomotion, segments’ contraction rates, durations, and timing relationships changed in nonlinear trends from posterior to anterior. Movement features operating at various scales— segments, adjacent segments, and groups of segments—all varied along the body axis, each with a transition point around the mid-body. Moreover, distinct regions of the larval body axis displayed different degrees of flexibility in converting muscle recruitment to movement. These data support the model that posterior segments coordinate to power the visceral piston phase while anterior segments tolerate greater flexibility to enable bending and turning. In short, our findings reveal an unanticipated degree of axial heterogeneity in the coordination of limbless crawling, opening new avenues to study the origins of whole-body coordination and the nature and consequences of segmental diversity.

## MATERIALS AND METHODS

### Animal care and genotypes

*Drosophila melanogaster* strains used were the following:

w; +/+; Mef2-GAL4/TM6B-Dfd-YFP (derived from RRID:BDSC_27390)

w; +/+; UAS-jGCaMP7f (RRID:BDSC_79032)

w; +/+; Mef2-GAL4, UAS-Gerry/TM6, Tb, TubP-GAL80 (derived from RRID:BDSC_27390 and RRID:BDSC_80141)

*Drosophila* stocks were maintained at 25° C on molasses-based fly food. Fly crosses and embryo/larval collections were maintained on yeasted apple juice agar plates at 25° C, humidified to at least 75%.

#### Obtaining larvae for imaging

Larvae of genotype *w; +/+; Mef2-GAL4,UAS-Gerry/Mef2-GAL4,UAS-Gerry* (“Gerry”) were obtained from in-crosses of same-genotype adults. Larvae were screened for presence of GCaMP in body wall muscles prior to imaging sessions.

Larvae of genotype *w; +/+; Mef2-GAL4/UAS-jGCaMP7f* (“jGC7f”) were obtained from adult *w; +/+; UAS-jGCaMP7f* crossed to *w; +/+; Mef2-GAL4/TM6B-Dfd-YFP*. Larvae were screened for presence of GCaMP in body wall muscles prior to imaging sessions.

Embryos were obtained from crosses during a 24-hour collection window, aged for an additional 24 to 48 hours, and larvae to be imaged were subsequently screened and selected by size (average 1mm after 24 hours for first-instar jGC7f larvae; 2.5mm after 48 hours for second-instar Gerry larvae). Larval staging was confirmed by examining larval mouth hooks under magnification to check for characteristic second-instar size and shape. Larvae of both sexes were used and sex was not determined, as sexual characteristics have not yet emerged at these instars. Larvae were handled using soft-bristled paintbrushes.

### Imaging larval crawling inside agarose channels

For in-depth explanation of steps, including recipes used, see Greaney and Heckscher, 2024.

#### Channel preparation

Linear agarose channels were cast with 2% (w/v, in distilled water) LE agarose (APExBIO, Houston, TX) against raised channel molds (Heckscher et al., 2012; Greaney and Heckscher, 2024). Channels used for first-instar jGC7f larvae were either 200 µm (n=14 larvae) or 250 µm (n=4 larvae) wide by 200 µm deep; channels used for second-instar Gerry larvae were approximately 450 µm wide by 450 µm deep. After the agarose fully set, channels were unmolded, trimmed to fit on slides, and kept at 4° C immersed in water until 15 minutes prior to use. Prior to use the channels were transferred to room temperature. Larvae were matched to channels of appropriate widths relative to body size: larvae were snugly enclosed by channel walls and coverslip, which reduced the speed of crawling (facilitating tracking) and prevented larvae from turning around inside channels.

#### Imaging larval locomotion

After screening larvae for expression of the appropriate fluorophore, larvae were kept on unyeasted apple juice agar plates for the duration of an imaging session (average 2 hours). We observed no effect of time within session on animals’ crawling speeds, periods, or stride distances, and therefore grouped all animals together for analysis. Prior to imaging, larvae were washed in distilled water, then placed into an agarose channel, immersed in distilled water, and covered with a glass coverslip (Figure 1A). Channels were saturated with water throughout the duration of video capture.

**Figure 1:**
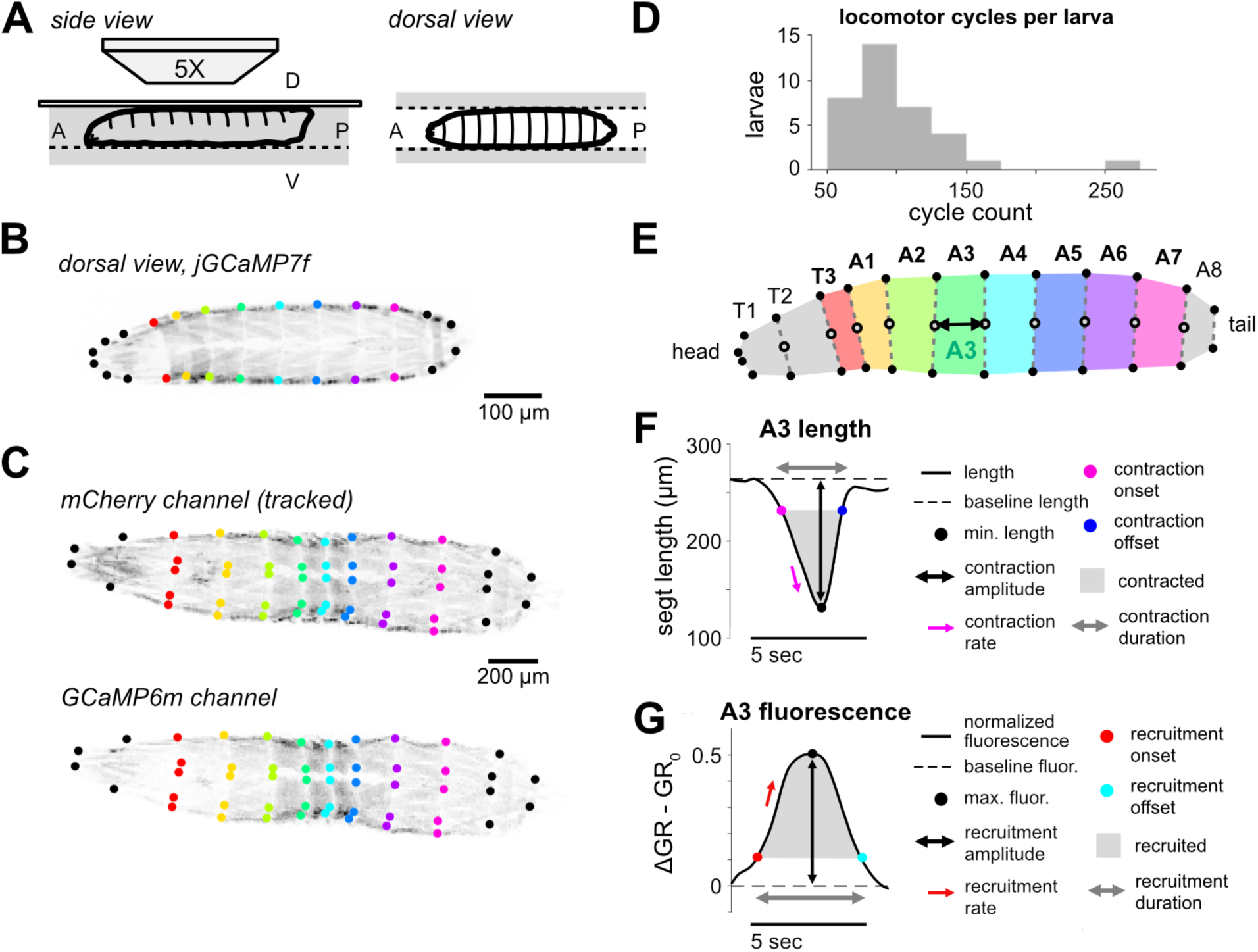
Segment-level kinematic and muscle recruitment data were obtained for many cycles of larval locomotion. A: Schematic of confocal imaging setup. B: Example frame for a crawling jGCaMP7f larva. Grayscale values, jGCaMP7f fluorescence intensity (inverted). Colored dots, DeepLabCut keypoints at the edges of each segment boundary. C: Example frame for a crawling Gerry larva. Red (top) and green (bottom) emitted fluorescence were collected in separate channels. D: Locomotor cycles obtained per larva imaged, n=35 larvae. E: Schematic representation of larval position. Black dots, keypoints; white dots, segment boundary centroids; double-headed arrow, example segment length. F: Schematic of contraction features. G: Schematic of fluorescence features.

Larvae were imaged one at a time for 5-20 minutes. jGC7f larvae were imaged using a Marianas spinning disk confocal microscope (3i, Denver, CO) with 488nm laser illumination, at 20 Hz, using laser intensities that prevented range saturation. Gerry larvae were imaged using a Zeiss LSM 800 point-scanning confocal microscope (Zeiss, Oberkochen, Germany) with 488nm and 561nm laser illumination, at 13.3 Hz, using laser intensities that prevented saturation in either channel. All larvae were imaged using a 5X, 0.15 NA objective (Zeiss). To keep larvae in the field of view, motorized stage movements were commanded manually throughout the imaging session.

Larvae were allowed to crawl until the end of video capture (12k frames, for jGC7f larvae), until they paused crawling for a period of at least 20 seconds, or until they reached the end of the linear channel, whichever happened first. To acquire additional cycles, some larva were allowed to recover for 1 minute outside of the channels, then were replaced and recorded again. All videos recorded of a single larva were combined for analysis.

### Developing DeepLabCut (DLC) segment tracking pipeline

To track larval segment boundaries, the DeepLabCut (DLC) Python package was used (Mathis et al., 2018; RRID:SCR_021391). The following steps resulted in boundary positions over time for segments T3 through A7. Subsequent analysis was performed in Matlab version 2021b (RRID:SCR_001622).

#### Initial training

Videos were oriented so that the larva’s head pointed left. If larvae were rolled relative to the imaging plane, videos were flipped so that the more visible side was the larva’s left side.

Separate DLC networks were trained, each using different tracking keypoints, for videos of jGCaMP7f-expressing larvae (“GCaMP network”) and videos of Gerry-expressing larvae (“Gerry network”).

The GCaMP network tracked two keypoints per segment boundary (Figure 1B), as well as two keypoints each to estimate locations of segments T2 and T1 and two points to estimate the position of a pair of longitudinal tail muscles (dorsal external oblique 1; Campos-Ortega and Hartenstein, 1997). Keypoints were placed at the exterior-most visible point of the left and right sides of a segment boundary. In frames where an animal was rolled substantially to one side, the ventral-anterior corner of ‘muscle 4’/‘lateral longitudinal 1’ was labeled instead.

The Gerry network tracked five keypoints per segment anterior boundary (T3 through A7). Keypoints were placed at the exterior-most visible anterior edges of the left and right sides of each boundary; the central anterior corner of dorsal longitudinal muscles (dorsal oblique 1; Landgraf et al., 1997); and an additional point at the ventral-anterior corner of the left side’s dorsal acute 3 (Figure 1C). This network also tracked four keypoints for the anterior boundary of segment A8, two points on the tail segment’s dorsal external obliques, and two keypoints to estimate the anterior boundary of segment T2.

Training data included 33 of 35 larvae from the analyzed dataset, as well as data from 61 additional larvae that were not analyzed further due to few overall locomotor cycles, unstable roll angle with respect to the imaging plane, poor manual tracking, or too few segments in view at any given time. 3550 total training frames were used. In both networks, the boundary between segments A8 and A9 was not tracked, and the anterior boundaries of T2 and T1 could not be tracked reliably. Therefore, only segments T3 through A7 were included for subsequent analyses.

#### Refining DLC network performance

An iterative refinement process was used to add training frames and arrive at the final GCaMP and Gerry DLC networks. First, a DLC network was trained using an initial set of training frames (approximately 900 frames from 30 videos for either network), then used to predict keypoint locations for all videos (either GCaMP or Gerry) in the dataset. Next, prediction quality was evaluated using a multi-step process. To do so, keypoints were converted into segment lengths (see below), then a list of potential problem frames was generated by finding frame-to-frame keypoint jumps exceeding 5 pixels, and by visually inspecting segment lengths plots for obviously problematic frame-to-frame changes. These potential problem frames were validated manually. Keypoint locations were corrected for a representative selection of these frames (approximately 200-600 frames per iteration), then the network was retrained with the addition of the corrected frames. This process was repeated until all videos were predicted with few (∼15 or fewer) problem frames. Any remaining problem frames were corrected manually.

### Converting keypoints to kinematic parameters

#### Filtering, finding segment boundary positions and lengths

Keypoint positions were smoothed using a 2nd-order Savitzky-Golay filter with a window of 15 frames, then converted into segment boundary positions by taking the mean of all keypoint positions belonging to the same segment boundary as the ‘midpoint’ for that boundary. Segment lengths were defined as the Euclidean distance between the midpoints of a segment’s anterior and posterior boundaries (Figure 1E). A “baseline” length for each segment was defined as the 95th percentile of its length distribution over all frames.

To estimate how far a larva traveled on each wave, segment positions were converted from frame (pixel) coordinates into real spatial coordinates. For the Gerry videos, frame coordinate positions were converted to real spatial coordinates using recorded stage movement data. For the jGC7f videos, which were captured using software/hardware that did not allow for motorized stage logging, stage movement was estimated as follows. If 40% or more of non-contracting segments’ keypoints exceeded a threshold velocity of 0.3 pixels/frame on a given frame, the stage was defined as moving. (Thresholds were chosen to produce the best agreement between visual inspection of videos and frames that were algorithmically defined as moving.) Then, non-contracting segments’ keypoint velocities were ranked by their similarity. The top 40% of most-similar velocities were selected, and the median of this set was taken as the estimated stage velocity and used to convert frame coordinates to real coordinates.

Finally, segments’ boundary positions were extrapolated on frames where they temporarily left the field of view, as follows. First, the expected distance between that boundary and its nearest visible neighbor was defined as the median distance between them in the original dataset (prior to any velocity and position estimation). Then, each time a boundary (typically at the head or tail end of the body) went out of frame, its positions following the out-of-frame period were compared to the position of the adjacent, more central boundary. A single value was added or subtracted to all of the out-of-frame boundary’s positions after the out-of-frame period, such that the median distance between the two boundaries after the out-of-frame period was corrected to match their expected distance. This process was repeated following any block of missing frames. Importantly, extrapolated boundary positions were only used to determine body segment movement distances and cycle stride lengths; they did not affect the calculation of tail movement times, segment lengths, or segment contraction timings.

#### Segment and tail forward movement times

To define periods of forward movement for each boundary, first, its smoothed positions over time (in real spatial coordinates) were converted to velocities. Boundary forward movement periods were defined as times when a segment’s frame-to-frame displacement exceeded 1% of its baseline length (for jGC7f larvae; 1.5% for Gerry larvae). Since no baseline length could be reliably found for the tail, an empirically-determined threshold of 2 µm/frame was used as the tail forward movement threshold. This threshold produced good agreement between manual definition of tail movement times (based on visual inspection of videos) and algorithmically-defined tail movement times.

#### Segment movement distances

Having established each segment’s moving periods, a segment’s movement distance on a given cycle was calculated using its anterior boundary. First, the forward movement period was found that overlapped with the segment’s contraction during that cycle’s locomotor wave (see below). Then, the boundary’s median positions before and after the movement period were determined, using all ‘not moving’ frames since the preceding ‘moving’ period to determine before-moving position, and all ‘not moving’ frames until the following ‘moving’ period to determine the after-moving position. The difference between before-moving and after-moving positions was taken as the movement distance for that cycle.

#### Segment contraction times and lengths

Segments’ lengths traces were processed to determine contraction onset, peak, and offset times as follows.

First, contraction peaks (length minima) were isolated by inverting the lengths traces and applying Matlab’s findpeaks function. To avoid finding small length changes that did not correspond to the primary contraction during a wave, a minimum peak prominence was enforced of 20% of the difference between segment’s 90th and 10th percentile lengths (for jGC7f larvae; 30% for Gerry larvae).

To find contraction onset and offset times, each contraction peak was first isolated from the rest of the lengths trace by finding the length peaks on either side. Then, during this window, the contraction onset was defined as the linearly interpolated time when the length decreased to 75% of the difference between maximum length before contraction and the minimum length during contraction. Similarly, the contraction offset was similarly defined as the time when the length increased to 75% of the difference between the minimum length during contraction and the maximum length after contraction (Figure 1F). This method was found to be robust to variation over time in between-cycle segment length.

#### Forward locomotor waves

Forward locomotor waves were defined as tail-to-head progressions of segmental contractions, as follows. The start of a forward wave was defined as the time at which the tail began moving forward. After this time, a “completed” forward wave occurred if each segment from A7 to T3 contracted sequentially. To allow for the scenario in which neighboring segments began shortening at around the same time, up to five ‘out-of-order’ frames were permitted in which an anterior segment apparently contracted before, or on the same frame as, its posterior neighbor. No rule was applied that segments had to de-contract in order.

To ensure that only complete, reasonably comparable forward waves were analyzed, waves/cycles were excluded if they met any of the following criteria:

- Not all 8 segments (T3-A7) contracted.
- One or more segments contracted more than once.
- One or more segments remained contracted for longer than the full wave duration (cycle period, see below).
- One or more segments’ defined contraction phases was far too early or late to plausibly belong to the same cycle (cycle phase < -2 or > 2).
- The wave preceded or followed a long pause, or a long pause happened partway through the wave. A long pause was defined as 1 second elapsing without any segment boundary moving forward.
- A cycle period could not be defined due to the tail not being visible at the start of the wave.
- The head end (including at least T3 and A1) was retracted, including at the beginning or end of the wave. Head retractions were defined as periods in which one or more contiguous anterior segments (between T3-A3) contracted and de-contracted without simultaneous co-contraction of the next posterior segment.
- After applying earlier exclusions, any remaining cycle whose period exceeded 4 standard deviations away from the mean cycle period in either direction was excluded.

Results of applying the above quality control criteria are summarized in Table 1.

**Table 1:** Results of wave quality control procedure. For each genotype, left two columns show mean number of putative waves prior to and median number of waves after quality control (see also Figure 1D). Remaining columns provide range (and mean in parentheses) of putative waves discarded for the following criteria: occurred before/ after a long pause; included a head retraction; one/more segments had no or multiple contraction events; one/more segments contracted for longer than the cycle period; one/ more segments contracted at cycle phase < -2 or > 2; cycle period was > 4 standard deviations above average.

|  | mean waves found | median waves retained | pauses | head retractions | missing or multiple contractions | long contr. | bad contr. phases | bad cycle periods |
| --- | --- | --- | --- | --- | --- | --- | --- | --- |
| **jGC7f (n=18)** | 97.39 | 86.5 | 0–16<br>(3.56) | 0–8<br>(2.28) | 1–53<br>(10.72) | 0 (0) | 0 (0) | 0–1<br>(0.17) |
| **Gerry (n=17)** | 107.55 | 85 | 0–4<br>(1.05) | 0–10<br>(2.35) | 1–37<br>(8.45) | 0–11<br>(1.00) | 0–4<br>(0.4) | 0–1<br>(0.05) |

#### Cycle stride distance, period, and speed

Typically, to estimate forward stride distance for each locomotor wave, the visual centroid of a larva is tracked over time (Risse et al., 2017; Thane et al., 2023). To obtain an estimate that depended less on body perimeter estimation, stride distance for each cycle was estimated as the mean of distances moved by segments A2 through A7 on that cycle. Segments A1 and T3 were excluded from the estimation because they occasionally moved as part of a head movement between contractions, resulting in less reliable movement distance calculation for these two segments.

The start of each locomotor wave was defined (above) as the time at which the tail began moving forward in advance of segmental contractions. Cycle periods were defined as the duration from this tail movement until the following cycle’s tail movement.

Finally, each cycle was assigned an average speed by dividing the cycle stride distance by the cycle period.

#### Segment contraction phases

For each cycle, each segment’s contraction onset was converted into a cycle phase value. This was done by calculating the delay between the start of the wave and the segment’s contraction onset, then normalizing this delay by the cycle period.

### Extracting segment fluorescence traces

The calcium sensor GCaMP was used to obtain information about segments’ muscle recruitment on each cycle. Larvae expressed the Mef2-GAL4 driver, which drove expression of either jGCaMP7f (Dana et al., 2019) or Gerry (tethered GCaMP6m-mCherry, as in Ugur et al., 2017) in all body wall muscles. However, because of the high likelihood of movement artifacts during crawling, recruitment was only evaluated in Gerry-expressing larvae, in which dynamic calcium sensor (GCaMP) fluorescence could be normalized by stable fluorophore (mCherry) fluorescence.

#### Region of interest definition

To obtain calcium sensor activity over time (a “fluorescence trace”) for each segment, the DLC-tracked keypoints belonging to a segment’s anterior and posterior boundaries were connected to define the boundary around a region of interest. Then, all pixels fully contained within the region of interest were averaged. This produced a single fluorescence intensity value per frame, per segment. GCaMP and mCherry fluorescence were processed separately. If a segment was missing any keypoint from either boundary on a given frame, that frame was skipped.

#### Estimating green-to-red fluorescence ratios

To account for increases in GCaMP fluorescence due to movement or contraction rather than muscle recruitment, segment fluorescence traces were expressed as a change in green-to-red ratio over time (dGR - GR_0_). To obtain a baseline green-to-red ratio (GR_0_), the green-to-red fluorescence ratio (GR) was averaged across all frames during which a segment’s length exceeded its baseline length (i.e., the segment was unlikely to be recruited). A segment’s muscle recruitment was defined as the difference between GR and GR_0_ on each frame.

Fluorescence traces were Savitzky-Golay filtered, using the same kernel applied to segment lengths traces, prior to analysis. Missing data were not interpolated.

#### Controlling fluorescence data for viscera movement

For each segment, the middle 50% of pixels were defined by drawing lines passing through the 25% and 75% points of line segments connecting the anterior and posterior external keypoints. Any pixels falling fully or partially between these lines were excluded when calculating green and red fluorescence traces for the segment. Visual inspection confirmed that this removed most or all of visible gut fluorescence.

#### Motion correction of fluorescence data

As an additional safeguard against the possible effects of motion artifact, Two-channel Motion Artifact Correction (TMAC; Creamer et al. 2022) was used to infer segment recruitment using information about motion obtained from the red channel. For each video, green- and red-channel activity for each segment was isolated using DLC keypoints as described above, concatenated, and sent to TMAC as a set of 8 regions of interest. To avoid interpolating fluorescence data, only larvae for which all segments were visible throughout the entire video were processed (n=9 of 17 Gerry larvae). TMAC estimations of each segment’s activity were processed identically to fluorescence traces obtained using the green-to-red fluorescence ratio, and compared to these original results (only from the same 9 larvae), using the methods described next.

### Matching fluorescence and kinematic data (cycles)

Segments were processed one at a time to obtain the following recruitment metrics and when matching recruitment events into locomotor cycles:

#### Recruitment event timings and rates

To determine the peak, onset and offset timings and rates of segment recruitment events, fluorescence traces were treated identically to lengths traces, with the exception of a minimum peak prominence of 30% instead of 20%.

#### Matching recruitment events to cycles

Processing the trace as above led to one onset/offset time, one duration, and one rate value per detected recruitment event. Recruitment events were occasionally missing for a given segment’s contraction (due to missing keypoints), and some segments had more than one detected recruitment event on a given locomotor cycle. To find the recruitment events most likely to correspond to a segment’s contraction on each cycle, a ‘best match’ algorithm was used that paired a single recruitment event to each cycle. First, recruitment peak timings were compared to their segment’s length minimum timings, and each peak was provisionally ‘assigned’ to its closest matching minimum in time. Then, if there were any duplicate assignments (multiple recruitment peaks assigned to the same length minimum), only the nearest match was retained. After this process, the paired recruitment events (peaks, plus their associated onset/offset timings and rate values) were assigned to the same locomotor cycle as the matching contraction.

After matching, quality control steps were applied to discard any probable bad matches. First, if any recruitment onsets or offsets assigned to a given cycle occurred prior to an earlier cycle’s length minimum, or later than a later cycle’s length minimum, they were discarded. Applying a stronger assumption, if any recruitment onset, specifically, occurred later than the same cycle’s segment length minimum, all of that cycle’s matched recruitment events were discarded. Finally, if any delay between a segment’s recruitment onset and its contraction onset exceeded the median period of a full locomotor cycle, that cycle’s recruitment events were discarded. Thus, the remaining matched recruitment events occurred within a reasonable time frame relative to their assigned contraction events.

#### Recruitment phases

Having assigned recruitment events into cycles, phases were determined for these events using the same procedure as for contraction events: the delay between a cycle’s start time (tail movement) and a segment’s recruitment onset was normalized to the total cycle period.

### Accounting for cycle speed information

To investigate the extent to which intersegmental correlations arose from the relationships between segments’ contractions and locomotor speed, speed information was regressed away from segments’ kinematic data. To do so, separately for each segment, linear regression was performed to obtain a relationship between cycle speed and each of the segment’s contraction kinematic features (contraction amplitude, contraction rate, contraction duration). The predicted value of each feature on each cycle was then subtracted, leaving residual values. Pearson’s correlation coefficients were calculated between segments using these residual values in place of the original values.

### Modeling block correlation structure

To evaluate the spatial structure of contraction correlations among segments, a method was designed to find ‘blocks’ of contiguous segments that share a correlation strength. This method generates and compares block-based models of segments’ correlation strengths for a single feature of contraction kinematics (e.g., contraction amplitude).

#### Model structure and assumptions

The central assumption of the method is that, within a block, adjacent segment pairs share the same correlation strength for the kinematic feature in question. Then, all further pairwise correlations within the block are modeled as the expected correlation under ‘local-only’ correlations, which decrease exponentially as a function of distance. The correlation strength is determined for each block separately. Outside of blocks, segment pairs share a single baseline correlation strength (see example in Figure 10A).

In this method, blocks potentially range in size from including just one segment to all eight segments. Two blocks may overlap but blocks may not nest fully inside other blocks and a segment may not be contained in more than two blocks. The 8-block model corresponds to a null model in which each segment is uncoupled to any others, except for a constant average correlation strength for all segment pairs. Conversely, the 1-block model (see Figure 10B) corresponds to a null model in which correlation between all eight segments is purely dependent on distance (local-only), with no other variation. For all other models, in which subsets of segments group together inside blocks, expected correlation between segments is a function of whether or not they belong to the same block and, if applicable, the correlation strength for that block.

#### Defining and evaluating models

To fit a model given a set of block edges, the expected pairwise correlations were fit as follows (see Figure 10A).

- Within a block, we averaged the *d*’th root of all correlations, where *d* is the distance between the segments in the pair (e.g., for adjacent segments, *d*=1). To permit this operation, correlations less than zero were set to zero.
- When two blocks overlapped, we performed the same operation but took the *d*’th root of the correlation as being the average of the values in the two blocks. All within-block values could be estimated simultaneously in this way by linear regression.
- Out-of-block elements, which correspond to segment pairs that were not in the same block, were averaged.
- To prevent lower-than-expected long-range correlations within blocks, any within-block values lower than the out-of-block average value were raised to this value.

To find the most likely block structure for each larva’s empirical correlations, first, every possible model was fit (n=1094 possible block combinations). Then, the log likelihood of the data under each model was calculated assuming a Gaussian distribution with a covariance matrix equal to the fitted model. Finally, standard model comparison was performed using the Bayesian Information Criterion (Schwarz, 1978), which penalizes parameters to account for models with more parameters being likely to fit better. Results were similar when the Akaike Information Criterion (Akaike, 1974) was used for model selection instead.

The model as structured has one degeneracy: nearly-uncorrelated segments can be included in the same block with a very low fitted correlation, which model comparison considers more parsimonious than using several 1-segment blocks. To analyze results of block modeling, we therefore only evaluated ‘strongly-coupled’ blocks (r > 0.5 for segments with distance=1). However, when the same analyses were performed considering all blocks (Figures S7, S8), we found qualitatively similar results.

#### Alternative model assumptions

The method as described above assumes that correlations between segments within a block decrease with distance, until they reach a minimum (out-of-block average) value. A variant of the method was tested that assumed correlations did not decrease with distance (Figures S7, S8). In this version of the method, all segment pairs in a block were given the same correlation (the mean correlation strength of all in-block pairs from the empirical data). Out-of-block pairs were averaged as in the model above. Results from this model variant were similar. A second variant was tested in which correlation values within blocks were allowed to decrease with distance to values below the out-of-block average; results were again similar (results not shown).

### Experimental Design and Statistical Analysis

The number of cycles required per larva was estimated early in data collection, using the first five jGC7f larvae successfully predicted by a DLC network. For five individual larvae, a threshold of at least 50 ‘good’ forward locomotor cycles was determined by eye to produce reasonable error bounds around pairwise intersegmental correlations. Imaging sessions were designed to maximize the number of cycles that could be well-tracked and predicted per animal, and the minimum threshold of 50 cycles was maintained for the remaining imaged animals (see Figure 1D).

To compare distributions of kinematic or recruitment features across eight body segments in Figures 2 and 4–6, data were averaged across cycles for each larva. Distributions were compared using the nonparametric Friedman test and Holm–Bonferroni post-hoc multiple comparison correction applied to resulting p-values. To compare the same segment’s distributions of kinematic or recruitment features across data subsets or analysis methods in Figures 3, 4, and 7, data were again averaged across cycles for each larva, after which nonparametric Kolmogorov–Smirnov tests were used and Holm–Bonferroni post-hoc multiple comparison correction applied to resulting p-values. Corrected p-values for all pairwise comparisons performed are given in Tables 2–7.

**Figure 2:**
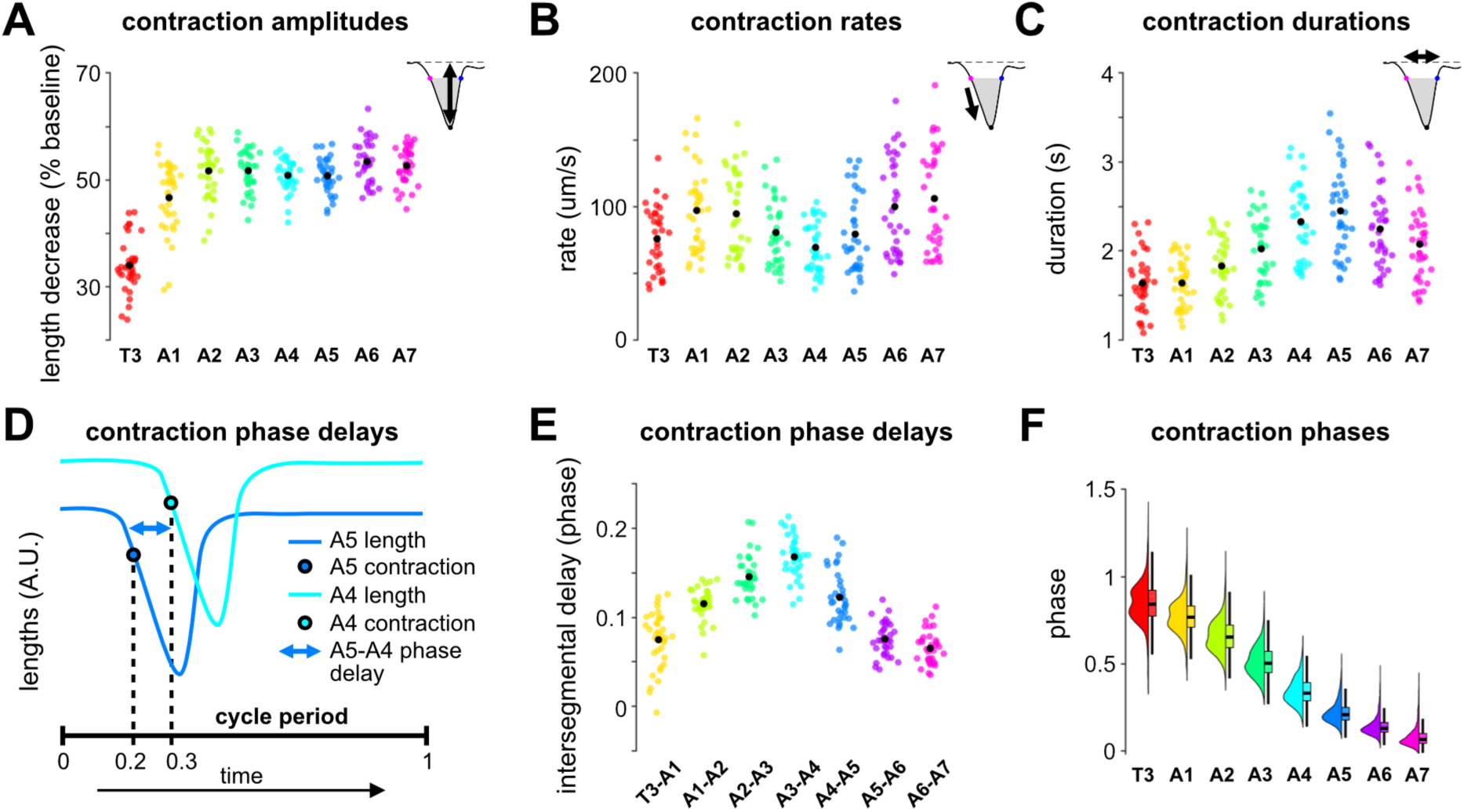
Contraction kinematics and phase relationships of individual body segments varied along the anterior-posterior body axis. A: Average normalized contraction amplitudes for segments T3 through A7 (black dots are overall means; n=35 larvae). Effect of segment was significant (nonparametric Friedman test: Q(279)=110.467, p=7.3 x 10^-21^). B: Average maximum shortening rates for T3 through A7. Effect of segment was significant (Q(279)=150.419, p=3.3 x 10^-29^). C: Average total contraction durations for T3 through A7. Effect of segment was significant (Q(279)=193.648, p=2.5 x 10^-38^). D: Schematic of calculating intersegmental contraction phase delays. Vertical offset between traces for clarity. For the simulated cycle shown, A5-A4 phase delay (black double-headed arrow) would be 0.1. E: Average intersegmental contraction phase delays between neighboring segments. Effect of segment pair was significant (Q(244)=153.086, p=1.7 x 10^-30^). F: Distributions of per-cycle contraction phases for segments T3 through A7. Histogram and box plot (without outliers) given for each segment. A–C, E: colored dots are individual larval averages (pooled across cycles); black dots are overall mean values; n=35 larvae. F: n=3497 cycles, 35 larvae. For pairwise comparisons among segmental distributions, see Table 2.

**Figure 3:**
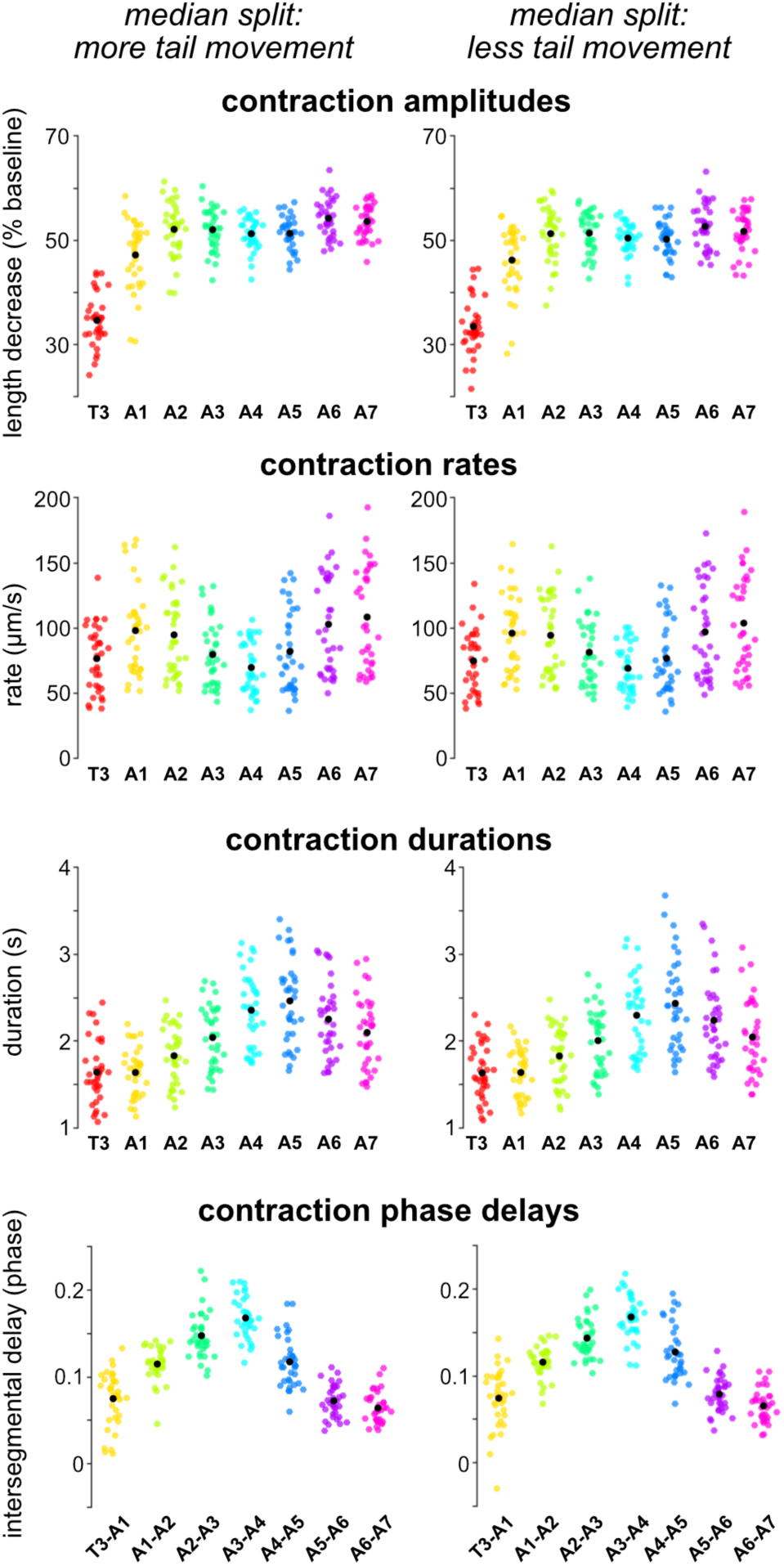
Axial trends in contraction kinematics did not depend on tail forward movement distance. Average values for the indicated feature of segmental contraction, pooled over the 50% of cycles where larvae moved their tails forward by a greater distance (left column) or by a lesser distance (right column). Median split of cycles was performed separately for each larva. Colored dots: individual larval averages; black dots: overall mean values; n=35 larvae. Kolmogorov–Smirnov tests with Holm–Bonferroni correction for multiple comparisons found no significant differences between greater-movement and lesser-movement distributions (see Table 3).

**Figure 4:**
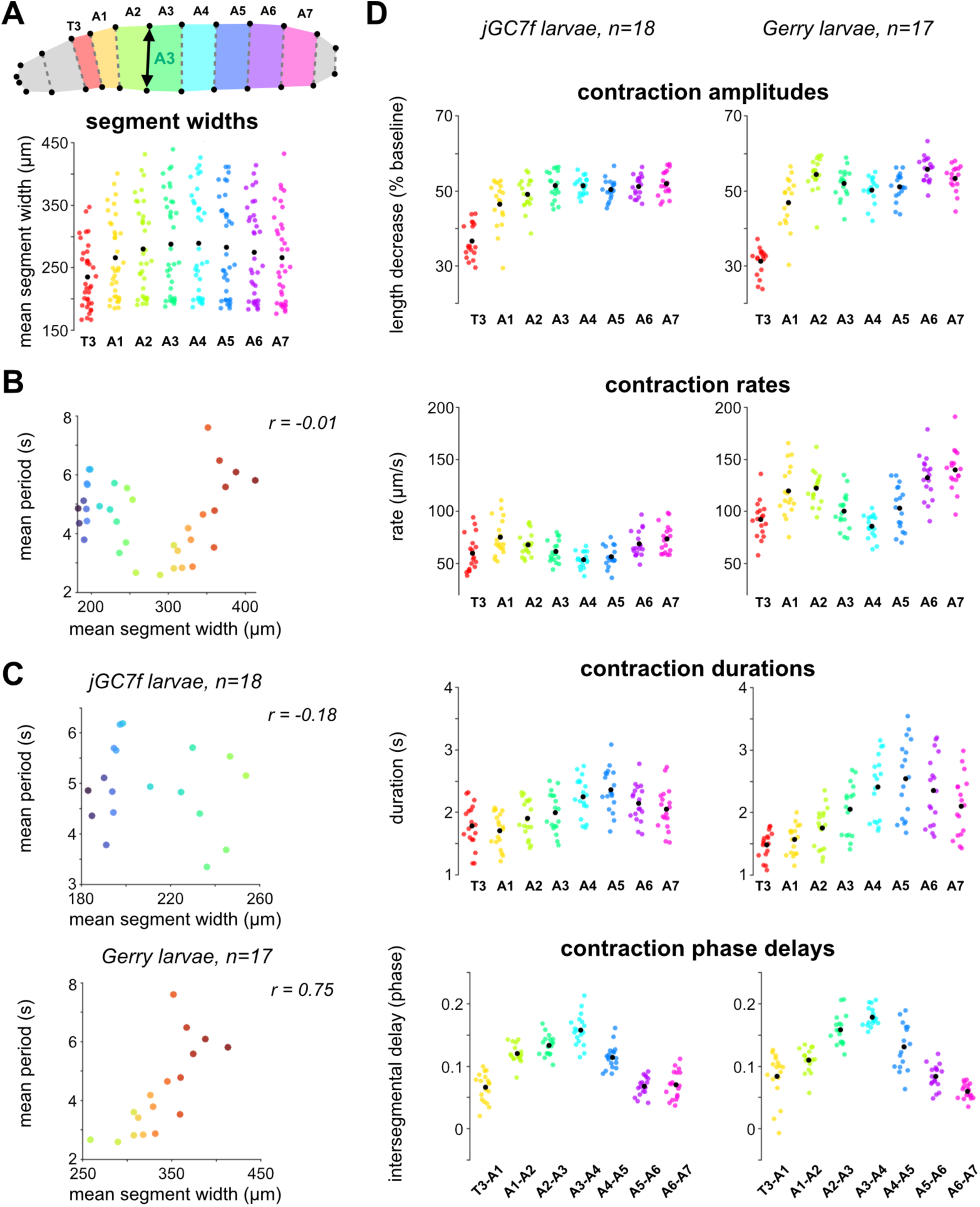
Axial trends in contraction kinematics are conserved across populations that appear differently constrained inside channels. A: (Top) Schematic of segment width calculation, which used the segment’s anterior boundary. (Bottom) Segmental distributions of mean boundary widths, n=35 larvae. Effect of segment was significant (nonparametric Friedman test: Q(279)=164.876, p=3.0 x 10^-32^). For pairwise comparisons among segmental distributions, see Table 4. B: Scatter plot of individual larvae’s mean segment widths against mean cycle periods. n=35 larvae; points colored per larva. C: Scatter plots of same data as in (B), separated into jGC7f (top) and Gerry (bottom) subpopulations. D: Average values for the indicated feature of segmental contraction, given separately for jGC7f (left column) and Gerry (right column) subpopulations. A, D: colored dots are individual larval averages (pooled across cycles); black dots are overall mean values; n=35 larvae. For Kolmogorov–Smirnov tests comparing jGC7f and Gerry distributions, see Table 4.

**Figure 5:**
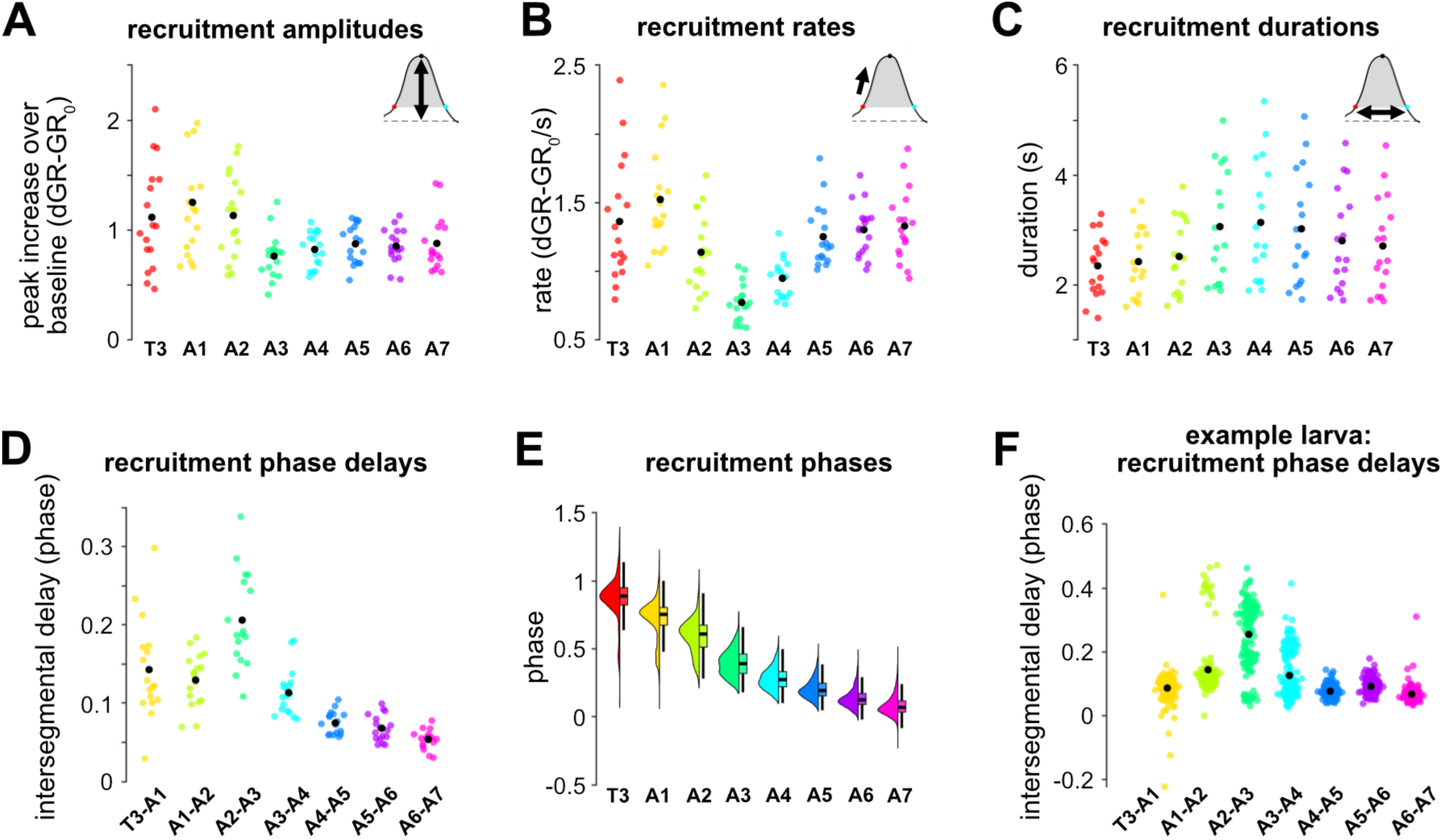
Recruitment and recruitment phase relationships of individual body segments varied along the anterior-posterior body axis. A: Average normalized recruitment amplitudes for segments T3 through A7 (black dots are overall means; n=35 larvae). Effect of segment was significant (nonparametric Friedman test: Q(135)=36.647, p=5.5 x 10^-6^). For pairwise comparisons among segmental distributions, see Table 5. B: Average recruitment rates (maximum rate of increase in ΔG:R-G:R_0_) for T3 through A7. Effect of segment was significant (Q(135)=72.098, p=5.6 x 10^-13^). C: Average recruitment durations for T3 through A7. Effect of segment was significant (Q(135)=89.412, p=1.6 x 10^-16^). D: Average intersegmental recruitment phase delays between neighboring segments. Effect of segment was significant (Q(118)=75.782, p=2.7 x 10^-14^). E: Distributions of per-cycle recruitment phases for segments T3 through A7. Histogram and box plot (without outliers) are given for each segment. Note that recruitment phases can be negative if (for example) segment A7 began increasing in fluorescence before the tail movement marking the start of a cycle began. F: Per-cycle intersegmental recruitment phase delays (colored dots) for a single example larva; n=155 cycles. Mean phase delay values given as black dots. A–D: colored dots are individual larval averages (pooled across cycles); black dots are overall mean values; n=35 larvae. For pairwise comparisons among segmental distributions, see Table 5. E: n=1633 cycles.

**Figure 6:**
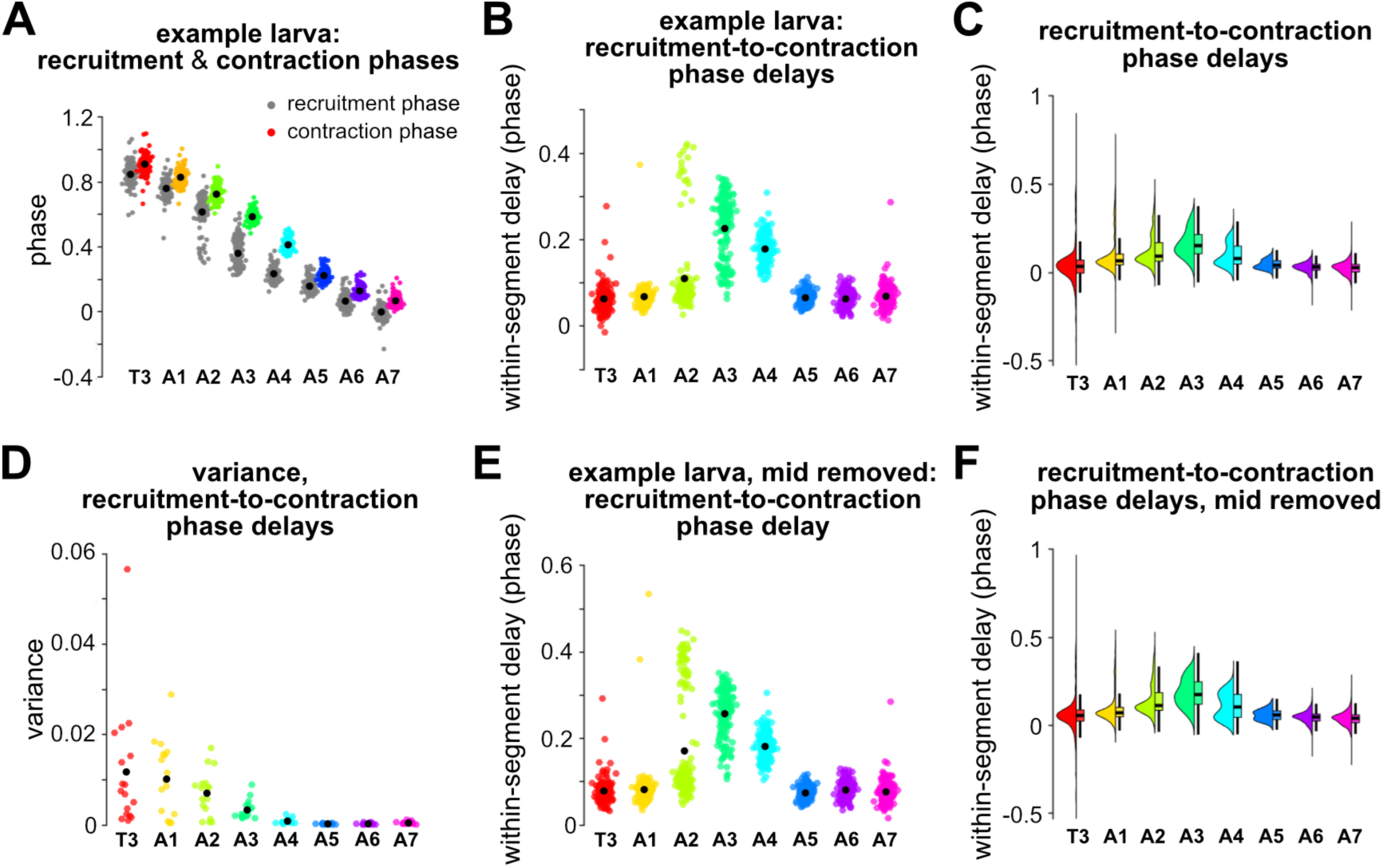
Phase delays between recruitment and contraction became longer and more variable in the mid-body. A: Segmental distributions of recruitment phases (grey dots) and contraction phases (colored dots) for all cycles from an example larva (“l21” in Figures S1, S2, S3). Black dots, means. B: Per-cycle phase delays between a segment’s recruitment and its contraction (colored dots) for the same example larva. Black dots, means. C: Segmental distributions of per-cycle phase delays between a segment’s recruitment and contraction, for all larvae. Histogram and box plot (without outliers) shown for each segment to highlight shape of distributions. Note that recruitment-to-contraction phase delays can be negative if a segment began shortening before substantially increasing in fluorescence. When values were pooled within-larva (n=17 larvae), effect of segment was significant (Q(135)=89.451, p=1.6 x 10^-16^). D: Average segmental variance in the phase delay between recruitment onset and contraction onset (black dots, overall mean variance values; n=17 larvae). E: Same as B, but after excluding midline 50% of pixels. F: Same as C, but after excluding midline 50% of pixels. C and F: n=1633 cycles across 17 larvae. For pairwise comparisons among segments and comparison of distributions in C and F, see Table 6.

**Figure 7:**
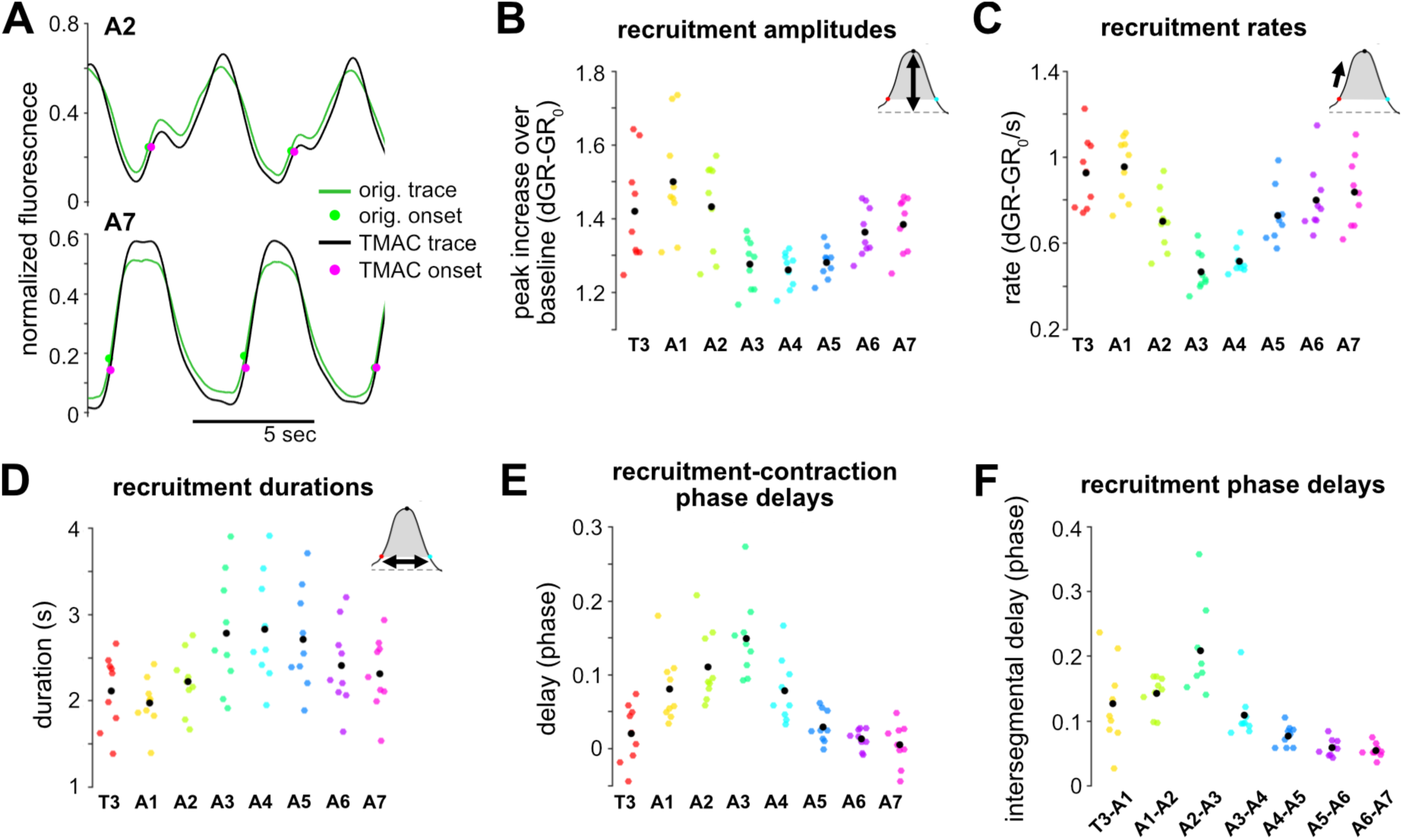
Axial trends in recruitment are conserved after algorithmic motion correction of videos. A: Comparison of original ratio (green) and ‘TMAC’ motion-corrected (black) fluorescence traces for segments A2 (top) and A7 (bottom) from an example larva during 15 seconds of video. TMAC traces represent inferred calcium sensor activity after correcting for bleaching and estimated red channel motion (Creamer et al. 2022). Fluorescence onsets found using original ratio traces: green dots; same onsets found using TMAC traces: magenta dots. Both original and TMAC traces normalized to minimum and maximum values during full recording (values between 0 and 1). B: Average segmental recruitment amplitudes, calculated using TMAC results. Effect of segment was significant (Q(71)=46.852, p=6.0 x 10^-8^). For pairwise comparisons among segmental distributions and Kolmogorov–Smirnov tests comparing TMAC results to original segmental distributions, see Table 7. C: Average segmental recruitment rates, calculated using TMAC results. Effect of segment was significant (Q(71)=43.259, p=3.0 x 10^-7^). D: Average segmental recruitment durations, calculated using TMAC results. Effect of segment was significant (Q(71)=52.852, p=4.0 x 10^-9^). E: Average phase delays between a segment’s recruitment and contraction, calculated using TMAC results. Effect of segment was significant (Q(71)=49.667, p=1.7 x 10^-8^). F: Average intersegmental recruitment phase delays between neighboring segments, calculated using TMAC results. Effect of segment was significant (nonparametric Friedman test: Q(62)=40.762, p=3.2 x 10^-7^). B–F: colored dots are individual larval averages (pooled across cycles); black dots are overall mean values; n=9 larvae without gaps in any segment’s visibility during recording period.

**Table 2:** Results of statistical tests accompanying Figure 2. Holm–Bonferroni adjusted p-values for Friedman’s tests comparing segments’ (or pairs of segments’) distributions of the indicated contraction feature (n=35 larvae). Bolded values: significant differences, alpha = 0.05. Note that p values >1 could occur due to the correction for multiple comparisons.

| **Friedman's test**<br>**(279) with Holm**<br>**correction** | 2A (cn.<br>amp.) | 2B (cn.<br>rate) | 2C (cn.<br>drn.) | **Friedman's test**<br>**(244) with Holm**<br>**correction** | 2E (cn. phase<br>del.) |
| --- | --- | --- | --- | --- | --- |
| **T3 vs A1** | 0.079 | **0.000** | 4.000 | **A1-T3 vs A2-A1** | **0.049** |
| **T3 vs A2** | **0.000** | **0.007** | 4.722 | **A1-T3 vs A3-A2** | **0.000** |
| **T3 vs A3** | **0.000** | 8.000 | **0.001** | **A1-T3 vs A4-A3** | **0.000** |
| **T3 vs A4** | **0.000** | 6.249 | **0.000** | **A1-T3 vs A5-A4** | **0.004** |
| **T3 vs A5** | **0.000** | 8.000 | **0.000** | **A1-T3 vs A6-A5** | 6.000 |
| **T3 vs A6** | **0.000** | **0.000** | **0.000** | **A1-T3 vs A7-A6** | 6.000 |
| **T3 vs A7** | **0.000** | **0.000** | **0.003** | **A2-A1 vs A3-A2** | 1.879 |
| **A1 vs A2** | 0.482 | 7.000 | 3.704 | **A2-A1 vs A4-A3** | **0.009** |
| **A1 vs A3** | 0.543 | **0.003** | **0.000** | **A2-A1 vs A5-A4** | 5.000 |
| **A1 vs A4** | 8.052 | **0.000** | **0.000** | **A2-A1 vs A6-A5** | **0.005** |
| **A1 vs A5** | 11.018 | **0.001** | **0.000** | **A2-A1 vs A7-A6** | **0.000** |
| **A1 vs A6** | **0.001** | 6.000 | **0.000** | **A3-A2 vs A4-A3** | 4.000 |
| **A1 vs A7** | **0.008** | 5.000 | **0.001** | **A3-A2 vs A5-A4** | 3.000 |
| **A2 vs A3** | 12.000 | 0.063 | 3.591 | **A3-A2 vs A6-A5** | **0.000** |
| **A2 vs A4** | 12.000 | **0.000** | **0.000** | **A3-A2 vs A7-A6** | **0.000** |
| **A2 vs A5** | 11.000 | **0.037** | **0.000** | **A4-A3 vs A5-A4** | **0.101** |
| **A2 vs A6** | 10.000 | 4.000 | **0.001** | **A4-A3 vs A6-A5** | **0.000** |
| **A2 vs A7** | 9.000 | 3.591 | 4.278 | **A4-A3 vs A7-A6** | **0.000** |
| **A3 vs A4** | 8.000 | 1.819 | 0.179 | **A5-A4 vs A6-A5** | **0.000** |
| **A3 vs A5** | 7.000 | 3.000 | **0.000** | **A5-A4 vs A7-A6** | **0.000** |
| **A3 vs A6** | 6.000 | **0.000** | 4.722 | **A6-A5 vs A7-A6** | 2.000 |
| **A3 vs A7** | 5.000 | **0.000** | 4.000 |  |  |
| **A4 vs A5** | 4.000 | 2.592 | 3.000 |  |  |
| **A4 vs A6** | 4.348 | **0.000** | 2.000 |  |  |
| **A4 vs A7** | 3.000 | **0.000** | 0.092 |  |  |
| **A5 vs A6** | 2.992 | **0.000** | 0.650 |  |  |
| **A5 vs A7** | 11.573 | **0.000** | **0.000** |  |  |
| **A6 vs A7** | 2.000 | 2.000 | 3.764 |  |  |

**Table 3:** Results of statistical tests accompanying Figure 3. Holm–Bonferroni adjusted p-values for Kolmogorov–Smirnov tests comparing high- vs. low-forward movement cycles for segments’ (or pairs of segments’) distributions of the indicated contraction feature. Bolded values: significant differences, alpha = 0.05.

| **Kolmogorov–Smirnov test with Holm correction** | 3A (cn. amp.) | 3B (cn. rate) | 3C (cn. drn.) | **Kolmogorov–Smirnov test with Holm correction** | 3E (cn. phase del.) |
| --- | --- | --- | --- | --- | --- |
| **T3** | 1.173 | 6.711 | 2.902 | **A1-T3** | 2.902 |
| **A1** | 2.561 | 1.934 | 5.872 | **A2-A1** | 3.355 |
| **A2** | 1.678 | 5.033 | 1.998 | **A3-A2** | 3.842 |
| **A3** | 1.921 | 2.902 | 5.872 | **A4-A3** | 1.998 |
| **A4** | 2.561 | 5.872 | 5.033 | **A5-A4** | 3.090 |
| **A5** | 2.207 | 4.194 | 4.836 | **A6-A5** | 3.842 |
| **A6** | 1.688 | 3.869 | 3.869 | **A7-A6** | 3.355 |
| **A7** | 0.750 | 6.711 | 5.122 |  |  |

**Table 4:** Results of statistical tests accompanying Figure 4. Holm–Bonferroni adjusted p-values. First results column: adjusted p-values for Friedman’s tests comparing segments’ widths distributions (n=35 larvae). Remaining results columns: adjusted p-values for Kolmogorov–Smirnov tests comparing jGC7f (n=18) and Gerry (n=17) larvae’s distributions of the indicated contraction feature. Bolded values: significant differences, alpha = 0.05.

| **Friedman's test (279) with Holm correction** | **4A (seg. wtdh.)** | **Kolmogorov–Smirnov test with Holm correction** | **4D (cn. amp.)** | **4D (cn. rate)** | **4D (cn. drn.)** | **Kolmogorov–Smirnov test with Holm correction** | **4D (cn. phase del.)** |
| --- | --- | --- | --- | --- | --- | --- | --- |
| **T3 vs A1** | **0.000** | **T3** | **0.038** | **0.000** | 0.125 | **A1-T3** | 0.121 |
| **T3 vs A2** | **0.000** | **A1** | 2.463 | **0.000** | 1.092 | **A2-A1** | 0.191 |
| **T3 vs A3** | **0.000** | **A2** | **0.004** | **0.000** | 1.092 | **A3-A2** | 0.121 |
| **T3 vs A4** | **0.000** | **A3** | 1.825 | **0.000** | 1.244 | **A4-A3** | **0.038** |
| **T3 vs A5** | **0.000** | **A4** | 2.463 | **0.000** | 0.935 | **A5-A4** | 0.191 |
| **T3 vs A6** | **0.003** | **A5** | 1.379 | **0.000** | 0.935 | **A6-A5** | 0.114 |
| **T3 vs A7** | 3.440 | **A6** | **0.049** | **0.000** | 0.496 | **A7-A6** | 0.231 |
| **A1 vs A2** | 0.111 | **A7** | 0.616 | **0.000** | 1.244 |  |  |
| **A1 vs A3** | 0.014 |  |  |  |  |  |  |
| **A1 vs A4** | **0.001** |  |  |  |  |  |  |
| **A1 vs A5** | 7.000 |  |  |  |  |  |  |
| **A1 vs A6** | 7.000 |  |  |  |  |  |  |
| **A1 vs A7** | 4.115 |  |  |  |  |  |  |
| **A2 vs A3** | 6.000 |  |  |  |  |  |  |
| **A2 vs A4** | 5.000 |  |  |  |  |  |  |
| **A2 vs A5** | 4.000 |  |  |  |  |  |  |
| **A2 vs A6** | **0.016** |  |  |  |  |  |  |
| **A2 vs A7** | **0.000** |  |  |  |  |  |  |
| **A3 vs A4** | 3.000 |  |  |  |  |  |  |
| **A3 vs A5** | 4.235 |  |  |  |  |  |  |
| **A3 vs A6** | **0.002** |  |  |  |  |  |  |
| **A3 vs A7** | **0.000** |  |  |  |  |  |  |
| **A4 vs A5** | 0.709 |  |  |  |  |  |  |
| **A4 vs A6** | **0.000** |  |  |  |  |  |  |
| **A4 vs A7** | **0.000** |  |  |  |  |  |  |
| **A5 vs A6** | 5.554 |  |  |  |  |  |  |
| **A5 vs A7** | **0.012** |  |  |  |  |  |  |
| **A6 vs A7** | 2.000 |  |  |  |  |  |  |

**Table 5:** Results of statistical tests accompanying Figure 5. Holm–Bonferroni adjusted p-values for Friedman’s tests comparing segments’ (or pairs of segments’) distributions of the indicated recruitment feature (n=17 Gerry larvae). Bolded values: significant differences, alpha = 0.05.

| Friedman's test (135) with Holm correction | 5A (rc. amp.) | 5B (rc. rate) | 5C (rc. drn.) | Friedman's test (118) with Holm correction | 5D (rc. phase del.) |
| --- | --- | --- | --- | --- | --- |
| T3 vs A1 | 17.989 | 6.391 | 11.000 | A1-T3 vs A2-A1 | 5.944 |
| T3 vs A2 | 17.000 | 14.000 | 11.000 | A1-T3 vs A3-A2 | 4.715 |
| T3 vs A3 | 0.086 | **0.003** | **0.000** | A1-T3 vs A4-A3 | 5.000 |
| T3 vs A4 | 2.639 | 0.645 | **0.000** | A1-T3 vs A5-A4 | 1.606 |
| T3 vs A5 | 16.000 | 14.000 | **0.000** | A1-T3 vs A6-A5 | 0.104 |
| T3 vs A6 | 15.000 | 13.000 | 2.000 | A1-T3 vs A7-A6 | **0.000** |
| T3 vs A7 | 14.000 | 12.000 | 10.000 | A2-A1 vs A3-A2 | 5.944 |
| A1 vs A2 | 13.000 | 0.092 | 9.000 | A2-A1 vs A4-A3 | 4.000 |
| A1 vs A3 | **0.001** | **0.000** | **0.000** | A2-A1 vs A5-A4 | 1.074 |
| A1 vs A4 | 0.086 | **0.000** | **0.000** | A2-A1 vs A6-A5 | 0.060 |
| A1 vs A5 | 3.142 | 8.762 | **0.000** | A2-A1 vs A7-A6 | **0.000** |
| A1 vs A6 | 4.369 | 11.000 | 1.376 | A3-A2 vs A4-A3 | 2.325 |
| A1 vs A7 | 3.718 | 10.000 | 8.000 | A3-A2 vs A5-A4 | **0.000** |
| A2 vs A3 | 0.067 | 0.645 | **0.011** | A3-A2 vs A6-A5 | **0.000** |
| A2 vs A4 | 2.201 | 9.000 | **0.000** | A3-A2 vs A7-A6 | **0.000** |
| A2 vs A5 | 12.000 | 8.000 | 0.079 | A4-A3 vs A5-A4 | 3.582 |
| A2 vs A6 | 11.000 | 7.000 | 7.000 | A4-A3 vs A6-A5 | 0.408 |
| A2 vs A7 | 10.000 | 6.000 | 6.000 | A4-A3 vs A7-A6 | **0.001** |
| A3 vs A4 | 9.000 | 5.000 | 5.000 | A5-A4 vs A6-A5 | 3.000 |
| A3 vs A5 | 8.000 | **0.002** | 4.000 | A5-A4 vs A7-A6 | 3.257 |
| A3 vs A6 | 15.946 | **0.000** | 2.840 | A6-A5 vs A7-A6 | 2.000 |
| A3 vs A7 | 17.989 | **0.000** | 0.047 |  |  |
| A4 vs A5 | 7.000 | 0.414 | 3.000 |  |  |
| A4 vs A6 | 6.000 | 0.092 | 0.221 |  |  |
| A4 vs A7 | 5.000 | 0.073 | **0.001** |  |  |
| A5 vs A6 | 4.000 | 4.000 | 10.071 |  |  |
| A5 vs A7 | 3.000 | 3.000 | 0.270 |  |  |
| **A6 vs A7** | 2.000 | 2.000 | 2.000 |  |  |

**Table 6:** Results of statistical tests accompanying Figure 6. Holm–Bonferroni adjusted p-values. First results column: adjusted p-values for Friedman’s tests comparing segments’ recruitment-to-contraction phase delay distributions (n=17 Gerry larvae). Second results column: adjusted p-values for Kolmogorov–Smirnov tests comparing Gerry (n=17) recruitment-to-contraction phase delay distributions, without (6C) vs. with (6F) the midline 50% of segment’s pixels masked out over time. Bolded values: significant differences, alpha = 0.05.

| **Friedman's test (135)**<br>**with Holm correction** | **6C (rc.-to-cn.**<br>**phase del.)** | **Kolmogorov–Smirnov test**<br>**with Holm correction** | **6F vs. 6C** |
| --- | --- | --- | --- |
| **T3 vs A1** | 0.253 | **T3** | 1.937 |
| **T3 vs A2** | **0.002** | **A1** | 2.324 |
| **T3 vs A3** | **0.000** | **A2** | 0.244 |
| **T3 vs A4** | 0.169 | **A3** | 2.018 |
| **T3 vs A5** | 10.993 | **A4** | 1.861 |
| **T3 vs A6** | 10.000 | **A5** | 2.018 |
| **T3 vs A7** | 9.000 | **A6** | 1.330 |
| **A1 vs A2** | 8.000 | **A7** | 2.324 |
| **A1 vs A3** | 9.123 |  |  |
| **A1 vs A4** | 7.000 |  |  |
| **A1 vs A5** | 9.123 |  |  |
| **A1 vs A6** | **0.025** |  |  |
| **A1 vs A7** | **0.011** |  |  |
| **A2 vs A3** | 6.000 |  |  |
| **A2 vs A4** | 5.000 |  |  |
| **A2 vs A5** | 0.208 |  |  |
| **A2 vs A6** | **0.000** |  |  |
| **A2 vs A7** | **0.000** |  |  |
| **A3 vs A4** | 10.993 |  |  |
| **A3 vs A5** | **0.005** |  |  |
| **A3 vs A6** | **0.000** |  |  |
| **A3 vs A7** | **0.000** |  |  |
| **A4 vs A5** | 6.778 |  |  |
| **A4 vs A6** | **0.014** |  |  |
| **A4 vs A7** | **0.006** |  |  |
| **A5 vs A6** | 4.000 |  |  |
| **A5 vs A7** | 3.000 |  |  |
| **A6 vs A7** | 2.000 |  |  |

**Table 7:** Results of statistical tests accompanying Figure 7. Holm–Bonferroni adjusted p-values for Kolmogorov–Smirnov tests comparing original recruitment results to results following Two-color Motion Artifact Correction (Creamer et al. 2022) for segments’ (or pairs of segments’) distributions of the indicated recruitment feature. n = 9 Gerry larvae with no missing frames. Bolded values: significant differences, alpha = 0.05.

| Kolmogorov–Smirnov test with Holm correction | 7B (rc. amp.) | 7C (rc. rate) | 7D (rc. durn.) | 7E (rc.-cn. phase del.) | Kolmogorov–Smirnov test with Holm correction | 7F (rc. phase del.) |
| --- | --- | --- | --- | --- | --- | --- |
| T3 | **0.010** | 0.078 | 3.015 | 1.915 | A1-T3 | 3.618 |
| A1 | **0.010** | **0.000** | 0.149 | 0.623 | A2-A1 | 2.872 |
| A2 | **0.007** | **0.037** | 1.750 | 0.623 | A3-A2 | 1.915 |
| A3 | **0.000** | **0.002** | 2.872 | 1.915 | A4-A3 | 3.618 |
| A4 | **0.000** | **0.000** | 2.872 | 2.412 | A5-A4 | 3.015 |
| A5 | **0.000** | **0.000** | 1.915 | 0.545 | A6-A5 | 2.872 |
| A6 | **0.000** | **0.002** | 1.750 | 0.467 | A7-A6 | 1.750 |
| A7 | **0.002** | **0.010** | 3.015 | 2.412 |  |  |

For significance testing of correlation strengths at multiple intersegmental distances (Figures 8, S6), all correlation values were first Fisher z-transformed to improve normality of the distributions. Then, two-tailed Student’s T-tests were used to compare correlations either to a null value of zero or against each other (speed-corrected vs. original correlations, Figure 8B, Figure S6B) and Holm–Bonferroni post-hoc multiple comparison correction was applied to resulting p-values. Corrected p-values are given in Table 8.

**Figure 8:**
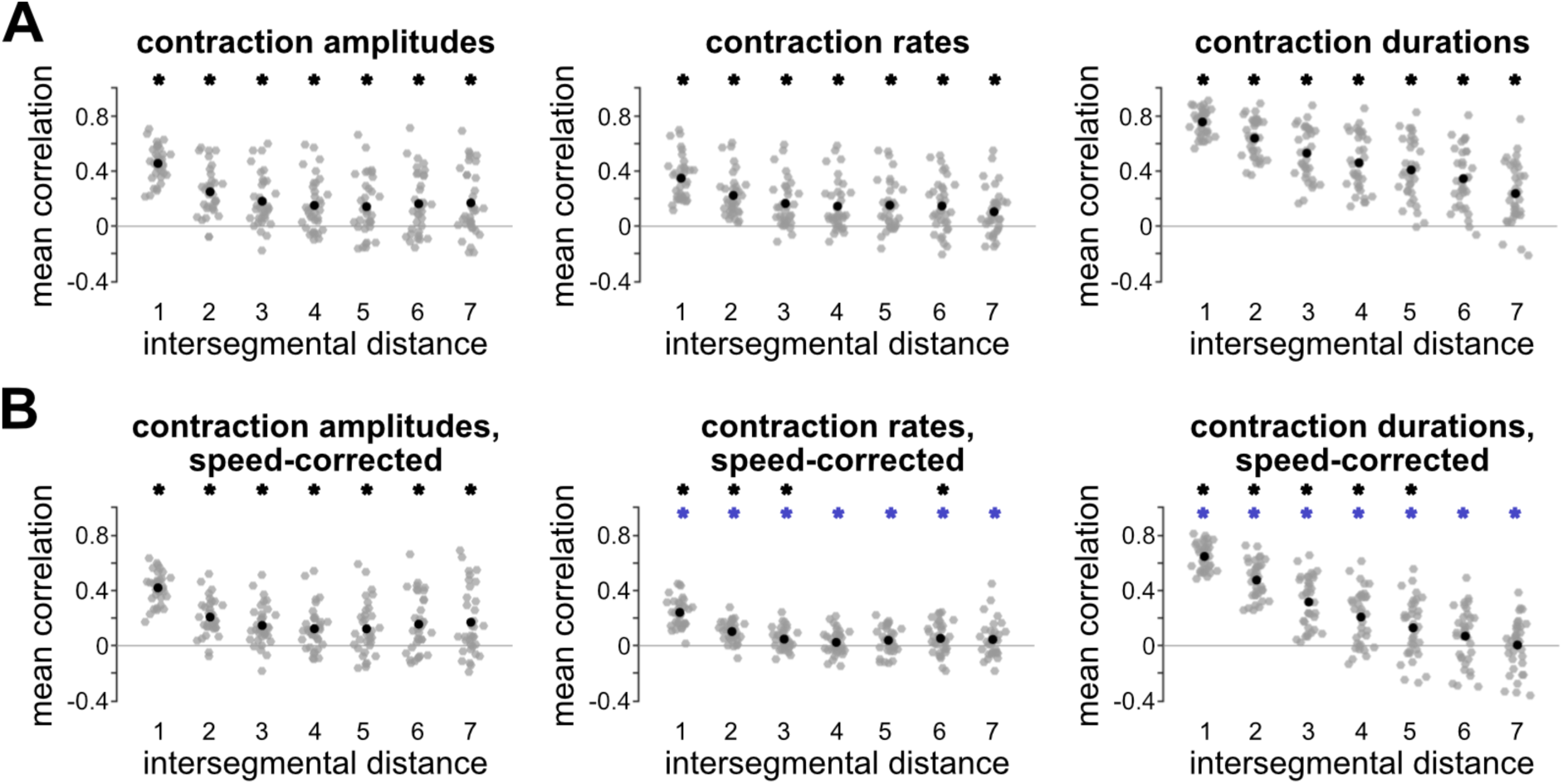
Contraction kinematics were correlated over long intersegmental ranges, with partial dependence on crawling speed. A: Pairwise intersegmental correlations of the indicated kinematic feature, averaged over all segment pairs at each intersegmental distance. Distance=1 indicates neighboring segments, etc. Asterisks: correlations significantly different from zero after Fisher’s z-transformation, two-tailed T-test, alpha=0.05. For p-values, see Table 8. B: Pairwise intersegmental correlations of speed-corrected residuals for each kinematic feature. To correct for speed, values for each segment were regressed against cycle speed and the result subtracted; correlations were calculated using residual values. Black asterisks: correlations significantly different from zero. Blue asterisks: correlations significantly different from original values (cf. panel A). For p-values, see Table 8. Grey dots: individual larval correlation values; black dots: overall mean values; n=35 larvae.

**Table 8:** Results of statistical tests accompanying Figure 8. Holm–Bonferroni adjusted p-values for two-tailed T-tests. Tests compared mean Fisher’s z-transformed correlation values at each intersegmental distance (n = 35 larvae) against a null of zero correlation (top and middle results sections) or between original and speed-corrected correlation values (bottom results section) for the indicated segmental contraction feature. Bolded values: significant differences, alpha = 0.05.

| **two-tailed T-test**<br>**(34)** | 8A (cn. amp. vs. 0) | 8A (cn. rate vs. 0) | 8A (cn. durn. vs. 0) |
| --- | --- | --- | --- |
| distance 1 | 0.000 | 0.000 | 0.000 |
| distance 2 | 0.000 | 0.000 | 0.000 |
| distance 3 | 0.000 | 0.000 | 0.000 |
| distance 4 | 0.000 | 0.000 | 0.000 |
| distance 5 | 0.000 | 0.000 | 0.000 |
| distance 6 | 0.000 | 0.000 | 0.000 |
| distance 7 | 0.000 | 0.001 | 0.000 |
| **two-tailed T-test**<br>**(34)** | 8B (cn. amp.,<br>speed-corr. vs. 0) | 8B (cn. rate,<br>speed-corr. vs. 0) | 8B (cn. durn.,<br>speed-corr. vs. 0) |
| distance 1 | 0.000 | 0.000 | 0.000 |
| distance 2 | 0.000 | 0.000 | 0.000 |
| distance 3 | 0.000 | 0.004 | 0.000 |
| distance 4 | 0.000 | 0.114 | 0.000 |
| distance 5 | 0.001 | 0.057 | 0.002 |
| distance 6 | 0.000 | 0.030 | 0.090 |
| distance 7 | 0.001 | 0.092 | 0.902 |
| **two-tailed T-test**<br>**(34)** | 8B (cn. amp.,<br>speed-corr. vs. orig) | 8B (cn.rate, speed-corr. vs. orig) | 8B (cn. durn.,<br>speed-corr. vs. orig) |
| distance 1 | 0.477 | 0.000 | 0.000 |
| distance 2 | 0.193 | 0.000 | 0.000 |
| distance 3 | 0.247 | 0.000 | 0.000 |
| distance 4 | 0.477 | 0.000 | 0.000 |
| distance 5 | 0.520 | 0.000 | 0.000 |
| distance 6 | 0.552 | 0.000 | 0.000 |
| distance 7 | 0.913 | 0.003 | 0.000 |

To distinguish observed correlations from the null model that correlation strength would depend on intersegmental distance (Figure 9C), bootstrapping was used to generate an expected distribution of correlation strengths under the null model. A separate distribution was generated for each larva. To do so, first, the larva’s cycles were sampled with replacement (up to the original number of cycles). Next, as for Figure 8, pairwise intersegmental correlations were calculated using the resampled data and Fisher z-transformed for normality. Then, the expected correlation at each intersegmental distance was calculated by averaging over all pairs of segments at a given distance (7 pairs of segments were averaged with distance=1, 6 pairs with distance=2, etc.). Lastly, the process was repeated (n=10,000 iterations). After generating bootstrapped distributions of expected correlation strength, empirical correlations (likewise Fisher-transformed) were compared to the bootstrapped distributions and a p-value calculated for each.

**Figure 9:**
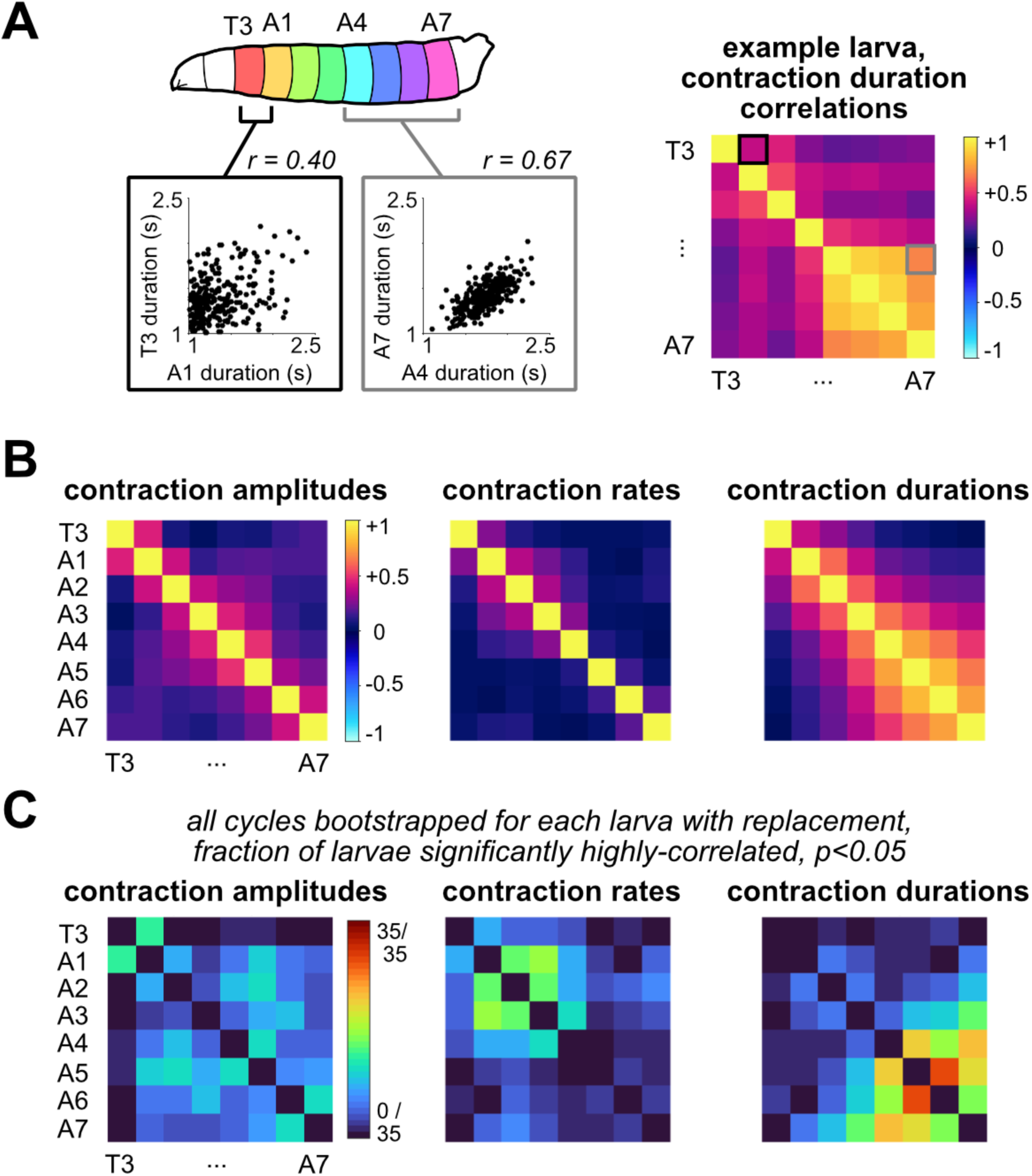
Range and strength of contraction duration correlations were higher among posterior segments. A: Schematic of calculating pairwise intersegmental correlation matrices. Example contraction duration relationships shown as scatter plots for two pairs of segments (left). Example plots correspond to two pairwise comparisons in the full correlation matrix (right), marked with boxes. B: Mean pairwise intersegmental correlations for the indicated feature of segmental contraction after speed correction (averaged over n=35 larvae). All values along the diagonal = +1 by construction. C: For each pair of segments, fraction of larvae with significantly (p<0.05) higher-than-expected pairwise correlations after speed correction (see Methods).

### Data and Code Accessibility

Experiments and analyses were not preregistered. Code will be made available at the following GitHub repository: https://github.com/kaufmanlab/larvariability-public. Raw video data and DLC annotations will be made available at the following FigShare repository: https://figshare.com/projects/Larval_locomotion_scales/272380.

## RESULTS

Our goal was to determine how segments are coordinated across the body axis during *Drosophila* larval visceral pistoning. We asked to what extent segmental coordination matches a model of uniform wave propagation through identical segments. We focused on three kinematic features of segmental movement during crawling—contraction amplitude, contraction rate, and contraction duration—as most directly impacting locomotion speed and therefore potentially needing to be coordinated across segments.

To study visceral pistoning specifically, we restricted larval behavior: In free crawling on flat surfaces, *Drosophila* perform a complex behavioral repertoire (Green et al., 1983; Kohsaka et al., 2017; Clark et al., 2018). To limit this repertoire to multiple consecutive cycles of visceral pistoning, we placed larvae inside linear channels (Heckscher et al., 2012; Greaney and Heckscher, 2024) (Figure 1A). To gain a broad understanding, we used various sizes (stages) and genotypes: 18 first-instar larvae expressing jGCaMP7f in all body wall muscles (Figure 1B; Movie 1) and 17 second-instar larvae expressing “Gerry,” GCaMP6m and mCherry, in all body wall muscle cells (Figure 1C; Movie 2). Distributions of larval sizes and cycle statistics (speeds, periods, and stride distances) are shown in Figure S1. We obtained an average of 100 forward cycles per animal (Figure 1D; Table 1). From all larvae, we identified segment boundary positions over time (see Methods; Movies 1 and 2), calculated the lengths of each segment from thoracic segment 3 (T3) to abdominal segment 7 (A7) over time (Figure 1E), and assessed the kinematic features of interest for each segment on each cycle (Figure 1F). From Gerry-expressing larvae, we also extracted each segment’s calcium fluorescence over time, a proxy for neuromuscular activity that we compare to kinematics (Figure 1G). Additional examples of segments’ contraction and fluorescence traces are shown in Figure S2.

### Segment contraction kinematics vary non-monotonically across the body axis

As a first step in testing a model of uniform segmental wave propagation during visceral pistoning, we analyzed the contraction kinematics of individual segments across the body axis. All three kinematic features—contraction amplitude, rate, and duration—varied non-monotonically. For amplitudes, the thoracic segment T3 contracted significantly less than the abdominal segments A2-A7 (Figure 2A, Table 2). For rates, shortening was slowest in the mid-body (A4 and A5; Figure 2B), as well as T3. For durations, posterior segments contracted for longer than anterior segments, with A5 contracting the longest (Figure 2C). Thus, segmental contraction waves do not uniformly propagate along the body. Instead, different segmental kinematic features vary in different non-monotonic patterns along the body axis.

### Propagation of the segmental contraction wave slows around mid-body

Coordinating the relative timings of specific movements is critical for behavior. In the case of visceral pistoning, the peristaltic wave depends on an ordered sequence of segmental contractions. The relative time at which each segment contracts can be expressed as a fraction of the total cycle period (i.e., its contraction phase, Figure 2D). The phase relationships between segments’ contractions may be linear, as in the uniformly propagating contraction wave during some types of swimming (Hill et al., 2003; Di Santo et al., 2021), or more variable, as in some types of terrestrial crawling (Stern-Tomlinson et al., 1986; Cacciatore et al. 2000, Trimmer and Issberner, 2007). Therefore, we asked whether *timing coordination* was uniform or variable along the body axis during visceral pistoning. We quantified the intersegmental phase delays between adjacent segments’ contractions (Figure 2D). Phase delays varied along the body axis and were slowest around mid-body (A4-A3; Figure 2E, Table 2), which appeared to result in a nonlinear distribution of segments’ contraction phases (Figure 2F). Thus, the segmental timing relationships for visceral pistoning in channels feature a slowing of the contraction wave around mid-body.

### Mid-body contraction wave slowing was not explained by tail movement or channel interactions

The axial variation in contraction wave propagation (Figure 2) could arise from several factors, including differences across segments in neural circuit activity and/or in conversion of muscle activation into forces via body-environment interactions (see, e.g., Peron et al., 2009; Srinivasan et al., 2012; Jonaitis et al., 2022; Kohsaka, 2023). Particularly because we constrained larvae inside channels as part of our approach, we wanted to evaluate the likelihood that physical forces such as body-channel interactions could explain the differences in wave propagation. Therefore, we performed two analyses to test for the influence of physical forces.

First, we tested whether differences in contraction kinematics could be explained by the forward movement of the tail. In forward visceral pistoning, the tail segments contract, pushing the viscera and center of mass forwards (Simon et al., 2010; Heckscher et al., 2012). We considered the possibility that waves with farther tail movement might slow at a more anterior point than waves with less tail movement. To test this, we split each larva’s cycles at the median into cycles with longer vs. shorter tail movement distances. We did not observe different axial trends for either contraction rates or durations (Figure 3, Table 3), arguing that the location of mid-body slowing was not a consequence of tail displacement.

A second possibility was that the axial differences in segmental coordination might directly result from physical interactions between the sides of the larva and walls of the channel, as larval segments were generally slightly wider in mid-body (Figure 4A, Table 4). Because the sizes of the larvae in our dataset varied, correlated with larval stage (Figure S1), we analyzed the first-instar and second-instar larvae separately. Our second-instar but not first-instar larvae appeared increasingly impeded by the channels at larger body sizes, leading to longer cycle periods (Figure 4B-C). Regardless of size and stage, however, we observed similar axial trends in the distributions of contraction amplitudes, rates, durations, and intersegmental phase delays (Figure 4D). Since both more- and less-constrained larvae shared the same non-monotonic patterns in segmental contractions, we conclude the structure of these distributions was unlikely to be a consequence of segments differing in their physical interactions with the channels.

### The relationship between recruitment and contraction waves becomes variable at mid-body

Since we could not explain the mid-body slowing of the contraction wave using proxies for physical forces, we next asked whether the non-uniform wave could be neural in origin. If so, similar non-monotonic axial trends would also be present in segments’ muscle activations. To assess segmental trends in neuromuscular activity, we quantified a proxy of muscle recruitment: segmental calcium activity in the Gerry-expressing larvae. We measured each segment’s recruitment amplitude, rate, and duration, which were closely analogous to the kinematic features, as well as phase delays in recruitment propagation.

First, we compared axial trends for recruitment and contraction features. Overall, trends for recruitment features were similar to analogous contraction features (compare Figure 2A-C to Figure 5A-C; Table 5). Further, the recruitment wave, like the contraction wave, slowed around mid-body, which appeared to result in a similarly nonlinear distribution of segments’ recruitment phases (compare Figure 2E-F to Figure 5D-E). Additionally, trends in the recruitment wave led similar trends in the contraction wave by one segment to the anterior. A spatial offset thus emerged between recruitment and contraction features: the slowest recruitment rates, longest durations, and longest phase delays each occurred one segment anterior relative to the slowest or longest analogous contraction feature.

Although the overall contours of axial trends were similar, unlike the contraction wave, the recruitment wave became strikingly variable in its propagation. Across larvae, this variability manifested particularly in the phase delays between A3 and A2 (Figure 5D); it was also visible within individual larvae as large cycle-to-cycle phase delay variability (example in Figure 5F). We conclude that, in addition to slowing around mid-body, recruitment wave propagation can progress differently on each crawl cycle, and the point of greatest variation in its progression appears to be in the anterior mid-body.

### The relationship between recruitment and contraction waves becomes variable at mid-body

Because, generally, we found high variance in the recruitment wave at mid-body (Figure 5D-F), we next investigated the timing relationships between the recruitment and contraction waves in single animals on a cycle-by-cycle basis. The relationship between a segment’s recruitment and its contraction can inform the role of muscle activations in locomotion. For instance, in anguilliform swimming, the phase delay between muscle recruitment and segment contraction changes systematically along the body axis, supporting different roles of muscle activity at the tail end and head end (Williams et al., 1989; Wardle et al., 1995; McMillen et al., 2008). Therefore, we directly asked whether any such systematic differences existed in larval crawling.

For each segment and each cycle, we calculated the phase delays between recruitment and contraction (Figure 6A-C). In most individual larvae, in at least one segment, the difference between recruitment and contraction phases was much larger and more variable than for other segments (example: Figure 6A-B, segments A2, A3; see also Figure S2C). Overall, the propagation of the recruitment wave was furthest in advance of the contraction wave at segment A3 (Figure 6C; Table 6). Moreover, the phase delays between recruitment and contraction, while consistent among posterior segments, became substantially more variable toward the head end, both within- and across larvae (Figure 6C-D), an effect that did not appear to depend on animal size (Figure S3). Interestingly, we observed relatively frequent negative phase delays between T3’s recruitment and contraction, indicating that shortening began before muscles had been substantially activated, possibly due to pushing from behind (Figure 6C, Figure S3).

One possible confound was the variable movement of viscera during visceral pistoning; because the viscera were visible in the fluorescence channels, this movement might have affected apparent segmental recruitment timings. To control for this possibility, we repeated this analysis while excluding the central 50% of each segment’s pixels (Methods). Results were unchanged (Figure 6E-F; Table 6), arguing that movement of visceral fluorescence was not the source of either the phase delay or the variability.

A further possible confound arises from how strongly larval body wall muscles deform during contraction, which makes it more likely that motion artifacts could be affecting changes in fluorescence over time. To reduce this possibility, we repeated our recruitment analyses using an alternate method of processing our red and green fluorescence channels: Two-color Motion Artifact Correction (TMAC; Creamer et al. 2022), which uses information from the red channel to model both motion and bleaching, thereby improving the estimation of calcium indicator activity. A comparison of the original ratio and TMAC-processed recruitment traces is shown for two example segments in Figure 7A. TMAC processing significantly affected the magnitudes of the inferred amplitudes and rates of segment recruitment (Figure 7B-C), but did not significantly affect segments’ recruitment timings or durations (Figure 7D-F, Table 7). Crucially, the overall axial trends identified earlier (Figure 5) were preserved nearly identically following TMAC processing. Thus, our results held when using a method of fluorescence processing that more actively accounted for potential motion artifacts.

In sum, delays in propagating recruitment and contraction between segments did not reliably align, leading to variable phase relationships between the recruitment and contraction waves beginning in the mid-body.

### Segments’ contractions are correlated over long ranges, only partially due to locomotor speed

Thus far, we have considered kinematic features on a segment-by-segment basis (Figures 2A-C, 5A-C) and their short-range adjacent pair relationships (Figures 2D-F, 5D-F). Short-range coordination might explain contraction kinematics during certain types of locomotion. However, a unique feature of visceral pistoning compared to other forms of locomotion is that head and tail movements are highly correlated as they are physically coupled to each other via the viscera and the fluid-filled body interior (Heckscher et al., 2012). Thus, for visceral pistoning locomotion, we must consider longer-range coordination. Below, we evaluated correlations between segments’ kinematic features over distance.

We determined which contraction features were significantly correlated and estimated the ranges, in intersegmental distance, over which correlations were observed. We found an average correlation strength for each feature at each intersegmental distance. For instance, at an intersegmental distance of three, the correlation was calculated by averaging values for the following segmental pairs: T3-A2, A1-A3, A2-A4, A3-A5, A4-A6, and A5-A7; whereas an intersegmental distance of eight was calculated using only the T3-A7 segmental pair (Figure 8A). At all intersegmental distances, all contraction features were significantly correlated (Table 8). Contraction durations were the most strongly correlated (Figure 8A), with average correlations being quite strong (r > 0.5) even up to a distance of three segments. We conclude that during visceral pistoning, segmental contractions are correlated over long ranges.

A somewhat trivial explanation for long-range correlations could be that they result from changes related to whole-animal locomotor speed. To account for this possibility, we generated a best-fit linear model for the relationship between speed and segments’ cycle-by-cycle contraction kinematics (Methods), subtracted this prediction from the data, and recalculated correlations using the residuals. Contraction amplitude correlations remained almost independent of speed (Figure 8B, Table 8). For contraction rates and durations, correlation strengths were significantly diminished, but for contraction durations, persisted at distances of up to five segments. This suggests that speed-related factors play a role in correlating certain features across the body, but cannot fully explain longer-distance correlations.

### Contraction and recruitment durations are strongly correlated among posterior segments

The above analysis revealed the existence of long-range coordination of segmental contractions, but did not localize correlations to a particular region of the body. Given that both segmental contraction and muscle recruitment varied across the body with a prominent mid-body transition, we asked whether long-range coordination also varied across the body, and if so, how (Figure 9A). In visceral pistoning locomotion, all segmental contractions move the exterior body wall; however, contraction of the posterior segments additionally pushes the interior viscera forward. The strength of correlations across groups of segments may reflect how tightly different body regions must be coordinated for behavior. Therefore, we wondered if coordination might differ among anterior or posterior groups of segments with different roles in behavior.

We performed the following analysis both with and without speed-corrected correlations, with similar outcomes (Figures S4, S5). First, for each kinematic feature—contraction amplitude, rate, and duration—we generated a correlation matrix with segments on each axis (Figure 9B; Figure S4A-C). Most strikingly, for contraction durations, correlations were much stronger in the posterior compared to the anterior (Figure 9B). We compared to a bootstrapped null model (Methods), finding correlations in segmental contraction duration were consistently higher than expected toward the tail end of the animal, at ranges spanning multiple segments (Figure 9C, right). Next, we repeated these analyses for muscle recruitment features. Generally, the correlations for muscle recruitment features were stronger than for analogous kinematic features (Figure S6), suggesting stronger coordination of segmental neuromuscular activity compared to segmental contractions.

Thus, independent of speed, contraction and muscle recruitment durations are coordinated more strongly and at longer ranges among posterior than among anterior segments.

### A new statistical model of segment interactions

Above, we observed that non-adjacent pairs of segments exhibited correlated contraction durations, and further that these correlations were more commonly elevated in the posterior segments. We hypothesized that there might be a strongly-coordinated posterior block of segments, whose contraction durations are adjusted as a unit from cycle to cycle, but weaker coordination among anterior segments, whose contraction durations could be more independently modulated.

To explicitly test the idea that groups of segments behaved as blocks during visceral pistoning, we developed a new statistical method. This method finds groups of segments with similar correlation strengths and identifies where along the body axis correlation strengths change. It compares actual data to all possible models in a model family, wherein segments are grouped into different numbers and locations of blocks of shared correlation strengths (Fig. 10A; Methods). The simplest model groups all segments in a single large block, which means correlation strengths do not change along the body axis (Figure 10B). More complex models group segments into two or more blocks, indicating that correlations change somewhere along the body axis. The best-fitting model was identified for each larva (Figure 10C); often, the best fit model had two strongly-correlated blocks of segments (Figure 10D). We thresholded the best-fitting model for each larva to retain blocks of segments with strong correlations (R > 0.5), and identified the segmental location of the blocks’ edges.

**Figure 10:**
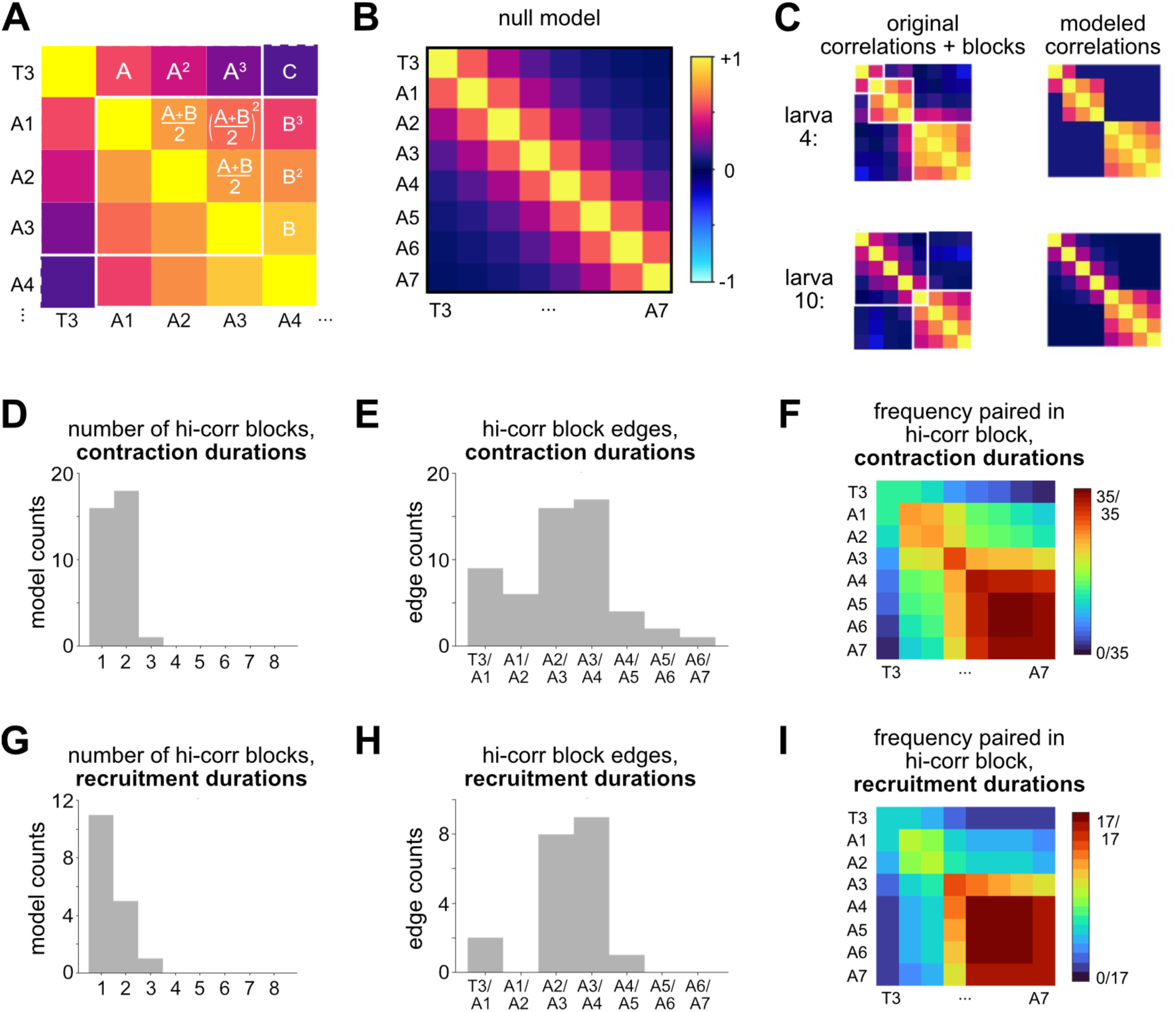
Contraction duration correlations were frequently higher among a group of posterior segments. A: Illustration of example model tested by the model-fitting procedure. Indicators “A” and “B” are adjacent-segment-pair correlation strength for different blocks; “C” is the out-of-block correlation strength. B: 1,094 total models were fit to each larva’s empirical correlation matrix, including the null model shown: all segments are included in the same block and correlation strength between segments is entirely a function of intersegmental distance. C: Two example best-performing block models (right) fit to empirical contraction duration correlation matrices for two individual larvae (left). After all possible block models are fit to a larva’s empirical correlation matrix, they are compared using Bayesian Information Criterion (BIC) and the best-scoring model is selected for further analysis. Color display scales and segment order as in B. D: Number of high-correlation blocks (adjacent-segment R>0.5) found by the best-performing block model for speed-corrected contraction durations for each of 35 larvae. E: Location of the edges of high-correlation blocks found by the best-performing models (n=35), indicating how often blocks ended between a given pair of segments. Edges between overlapping, similar-strength blocks were excluded (overlapping blocks with <0.2 difference in adjacent-segment R). F: Frequency with which best-performing models (n=35) included each pair of segments inside the same high-correlation block. The diagonal shows the frequency with which each individual segment was included in a high-correlation block, while the ‘exterior corner’ (T3 and A7 included in the same block) is equivalent to how many larvae’s correlations were best modeled as a single large block: the null model (as in B). G–I: analogous to D–F for recruitment metrics.

Block edges were infrequent between posterior segments A4-A7 (Figure 10E), suggesting that correlation strengths rarely changed between these segments. To check this more directly, we asked for how many larvae did the best-fitting models combine each pair of segments into the same block (Figure 10F). Posterior segments A4 through A7 were included in the same block with the other posterior segments for almost every larva (Figure 10F). Anterior segments, with the exception of A1 and A2, less often appeared together in the same block. The same observations held with and without accounting for speed or using variants of the model assumptions (Figure S7). In contrast to contraction durations, modeling blocks for contraction amplitude or rate correlations did not reveal consistent groups of segments (Figure S8). We repeated this analysis for muscle recruitment durations. The block results for recruitment duration correlations were strongly similar to those for contraction duration correlations (Figure 10G-I). Thus, generally, posterior segments comprised a coordinated block, and less often anterior segments also comprised a coordinated block.

One possible explanation for anterior-posterior differences in contraction feature correlation strength could be mechanical coupling promoted by interactions between segments and channel walls during crawling. For instance, in larvae that were more tightly constrained, the increased resistance to movement might make segments’ contraction and decontraction timings more interdependent than usual, potentially forcing groups of segments into acting as a more coordinated block; or, might introduce stochastic stick-slip interactions that instead decouple them. Therefore, we evaluated the relationship between larval width and contraction duration correlation strengths (Figure S9). Specifically, we calculated the ratio of posterior- vs. anterior-segment contraction duration correlation strengths, such that a larger ratio represented a larger difference across the body axis. Overall, the ratio decreased as larval width increased (Figure S9); and the effect was stronger for Gerry larvae, which appeared more impeded at increasing larval size (see Figure 3C). Thus, it is possible that physical confinement weakens the axial differences we observed.

## DISCUSSION

We determined how segments contract and how muscles are recruited during larval *Drosophila* visceral pistoning inside linear channels, finding previously unknown diversity in coordination across the body. First, segment-by-segment, muscle recruitment and contraction kinematics had their longest durations around the mid-body. Second, for adjacent pairs of segments, the propagation of contraction and recruitment waves slowed around the mid-body. Also around mid-body, the recruitment wave’s propagation became quite variable, and phase relationships between contraction and recruitment waves became inconsistent. Lastly, within a group of posterior segments, contraction and recruitment durations were strongly correlated. In summary, we discovered that features of visceral pistoning operating at various scales—segments, adjacent segments, and groups of segments—all vary along the body axis, each with a transition point around the mid-body.

For visceral pistoning locomotion, mid-body variation was unexpected. Compared to our study, previous studies of larval locomotion on flat surfaces sampled fewer cycles of movement (Gjorgjieva et al., 2013; Pulver et al., 2015; Sun et al., 2022; Liu et al., 2023) and did not find differences in segments’ contraction durations or amplitudes. Thus, the prevailing model has been uniform propagation of contraction waves along the body at all speeds (Gjorgjieva et al., 2013; Pehlevan et al., 2016; Liu et al., 2023). However, differences between the intrinsic properties, inputs, and input responses of anterior and posterior body segments are known (Srinivasan et al., 2012; Takagi et al., 2017; Tastekin et al., 2018; Murawski et al., 2020; Jonaitis et al., 2022; Jonaitis et al., 2024), motivating us to reevaluate this model.

Our study used a highly controlled preparation in which animals are confined within water-saturated agarose channels. This preparation offers clear advantages for imaging and quantification, enabling the collection of large numbers of similar visceral pistoning cycles and leading to our discovery of multiple scales of coordination along the body axis. However, locomotion could systematically differ in a channel environment compared to other contexts, including the flat surfaces often used to study *Drosophila* larval locomotion (Green et al., 1983; Dorgan, 2010; Vaadia et al., 2019; Booth et al., 2024). Thus, we cannot be sure that multi-scale coordination will be a prominent feature of *Drosophila* larval locomotion in all environments, for multiple reasons. First, friction will differ in channels compared to crawling over a flat substrate. Crawling over a substrate uses the ventral body’s traction against the substrate, whereas, inside a channel, friction is distributed across any points in contact with the channel (Dorgan 2010); if friction is irregularly distributed along the body axis, it might introduce irregular phase relationships across segments. Second, linear channels prevent some freedoms of movement, such as bending, which might in principle reduce expression of flexible coordination in the posterior. Third, being surrounded by water could introduce long-range hydrodynamic coupling of body segments, either through forces transmitted externally through the surrounding fluid or by amplifying the effects of internal fluid movement via external resistance. Finally, the form of locomotion itself may differ. As with leech and lamprey (Friesen and Kristan Jr., 2007; Islam and Zelenin, 2008), a different sensory environment could cause larvae to select a different motor output, such as tunneling or swimming. Therefore, we cannot rule out the possibility that larvae generate segmental recruitment and contraction relationships particular to the channel environment. While we did not find evidence that the degree of constraint affected overall patterns of segmental contraction (Figure 4), we did find it was weakly anti-correlated with strength of posterior-segment coordination (Figure S9), indicating that interactions with the environment affected at least some forms of coordination we measured. Imaging technologies are emerging that allow simultaneous recording of neural/muscular recruitment and segmental kinematics while larvae move on flat surfaces (Karagyozov et al., 2018; Bonnard et al., 2022; McNulty et al., 2025). These advances will enable testing how segmental coordination observed in channels changes when larvae navigate and perform more complex behaviors across different environments, and will contribute insight into the role of sensory feedback in diverse contexts (Hughes and Thomas, 2007).

The timing of muscle recruitment relative to the contraction wave varied along the body, with a tight relationship near the tail end that became quite variable starting in mid-body segments (Figure 6, Figures S2C, S3). This variability in muscle recruitment could signal that anterior segments are sensitive to sensory feedback about mechanical or internal state and adjust tension generated in advance of the contraction wave. Alternatively, the variability may be intrinsic to central circuits: recent work in nerve cord preparations showed variable propagation of neural activity through anterior segments, in the absence of sensory feedback (Francis et al., 2024). The variable delay between muscle recruitment and segment contraction highlights that converting neuromuscular activity to shortening is not a segment-autonomous process; other factors contribute to the conversion. These could include features of the channel environment (e.g., interaction with the channel walls) and dynamic (e.g., neuromuscular) or static (e.g., tissue mechanics) properties of the larval body. To test these possibilities, it will be essential to determine the mechanical forces that relate larval muscle activity to movement in a specific environment (Tytell et al., 2011; Ormerod et al., 2022; Kohsaka, 2023; Booth et al., 2024).

Additionally, we found differences in the strength of correlations for both contraction and recruitment durations among posterior and anterior segments. During visceral pistoning, we observed stronger correlations among a block of posterior segments, even when accounting for cycle speed (Figures 9, 10). Stronger correlations among groups of segments point to areas of the body that might need to be coordinated more tightly for behavior. An attractive possibility is that stronger posterior correlations may be due to differing behavioral roles for posterior and anterior segments. In the most posterior segments, contraction has a dual function: to move the exterior body wall and to push the internal viscera. The dual function of the posterior may require a tightly coordinated contraction; indeed, coordinated posterior activity, or inhibition of activity, has been noted even in the absence of locomotor waves (Pulver et al., 2015; Tastekin et al., 2018; Jonaitis et al., 2022). Additionally, during visceral pistoning, forward movements of the head and tail happen roughly synchronously with the contractions of multiple posterior segments (Movies 1, 2; Heckscher et al., 2012). Thus, stronger correlations within the posterior block may be partly explained by posterior segments’ individual correlations with head movements on a per-cycle basis; this possibility could be explored with improved methods for tracking head segments. By contrast, during visceral pistoning, anterior segments primarily move the exterior body wall and can be recruited between waves for lateral head movements (Lahiri et al., 2011; Berni et al., 2012; Gershow et al., 2012; Gomez-Marin and Louis, 2014; Wystrach et al., 2016). Therefore, the larva’s anterior segments may benefit from greater flexibility to respond to sensory inputs, enabling reorienting maneuvers.

What is the contribution of development to axial differences in locomotor coordination? Axial trends partially correlate with developmental factors. For example, larval thoracic and abdominal regions emerge from embryonic domains that differ in Hox gene expression (Roberts, 1971; Campos-Ortega and Hartenstein, 1997; Diao et al., 2024; Vasudevan et al., 2024). Manipulation of Hox genes leads to neural and behavioral differences in larvae (Dixit et al., 2007). We found that thoracic segment T3 contracted less tightly than abdominal segments (Figure 2) and sometimes shortened before muscles’ recruitment (Figure 6C, Figure S3), suggesting T3’s movements are subject to distinctive neural control. Furthermore, within the abdomen, the posterior-most Hox gene Abdominal-B can be expressed as far anterior as embryonic parasegment 10 (Delorenzi and Bienz, 1990), which gives rise to the border of larval segments A4 and A5. Posterior segments had stronger correlations among neuromuscular activity durations (Figure 10). Therefore, an attractive hypothesis posits segments with differing Hox codes feature differences in neuromuscular circuits that alter their timing relationships during a wave. Within the abdomen, a variety of segmental boundaries have been proposed as decision or reversal points with respect to locomotor waves and behavioral decision-making (e.g., Berni, 2015; Takagi et al., 2017; Tastekin et al., 2018; Murawski et al., 2020; Jonaitis et al., 2022). In all, our results align with the hypothesis that intersegmental differences affect the coordination and propagation of locomotor waves. Thus, it becomes of greater interest to understand the extent to which locomotor circuits differ along the body axis and how these circuit differences support axial coordination.

The need for an accurate model of visceral pistoning is at least three-fold: It contributes to the field’s understanding of embodied behavior; it is a required precursor for dissecting underlying neural mechanisms; and it motivates further investigation into the development of axially diverse motor circuits. Our findings of complex variation in how segments are recruited, contracted, and correlated across the body axis, alongside varying relationships between muscle recruitment and kinematics, make visceral pistoning locomotion in *Drosophila* larvae a rich system for interrogating locomotor control. Combined with the power of *Drosophila* genetics, this model unlocks new possibilities for probing the scales, mechanisms, and behavioral outcomes of whole-body coordination in segmented nervous systems.

## AUTHOR CONTRIBUTIONS

All authors designed the study; MRG performed the experiments and analyses; MRG wrote the paper with assistance from ESH and MTK; all authors edited the paper; MTK designed the block modeling; MTK and ESH supervised all aspects of the project; ESH provided funding.

## CONFLICT OF INTEREST STATEMENT

The authors declare no competing financial interests.

## ACKNOWLEDGMENTS

We gratefully acknowledge the assistance of H. Grier in setting up motion tracking and C. Wreden in animal care as well as the University of Chicago’s Statistics Consulting Group for helpful discussion of block modeling. This work was supported by the Pritzker Fellowship in the Neurosciences (MRG); NIH-NINDS F31NS118835 (MRG), and R01NS105748 and R56NS134862 (ESH).

## MOVIES

**Movie 1:** DeepLabCut network tracking of two minutes from example jGC7f larva video (ID: “larva 1”). Green channel gamma-adjusted for visualization purposes and displayed in inverted greyscale. Colored dots, tracked keypoints. Video display rate: 20 fps (∼real time).

**Movie 2:** DeepLabCut network tracking of two minutes from example Gerry larva (ID: “larva 22”). Red channel only, gamma-adjusted for visualization purposes and displayed in greyscale. Colored dots, tracked keypoints. Video display rate: 13 fps (∼real time).

## Notes

https://figshare.com/projects/Larval_locomotion_scales/272380

https://doi.org/10.6084/m9.figshare.31537603

https://doi.org/10.6084/m9.figshare.31508029

https://doi.org/10.6084/m9.figshare.31510339

### Competing Interest Statement

The authors have declared no competing interest.

### Summary of Updates

Added results using alternative motion correction method (Figure 7); changed analysis method to pool cycles within larva (Figures 2-6); added analyses to test for an effect of larval width/constraint on coordination (Figure S9); added further examples of segment contraction and recruitment traces (Figure S2); revised Methods section for clarity; revised Discussion section; changes to statistical tests performed; added Tables to report p-values and statistical significance; updated Figure 4B/C and corrected errors in Figure 4 legend.

