## Supplemental Figure Legends for "Multiple scales of coordination along the body axis during *Drosophila* larval locomotion"

**Supplemental Figure 1: Distribution of estimated larval sizes and locomotor cycle parameters.**

A: Distribution of segment baseline lengths for each larva; larvae sorted by increasing median length. n=8 segments per larva (T3 through A7); black dots, means.

B: Distribution of cycle speeds for each larva, sorted by same order as in A.

C: Distribution of cycle stride distances for each larva, sorted by same order as in A.

D: Distribution of cycle periods for each larva, sorted by same order as in A.

All panels: n=35 larvae. Pink box surrounds the Gerry subset of larvae.

**Supplemental Figure 2: Example segment length and fluorescence traces.**

A: Example segment length traces for 20 seconds of recording, jGC7f larva 6. Top three subpanels: contraction onsets (magenta dots), troughs (black dots), and offsets (blue dots) are labeled for the indicated segment. Bottom subpanel: lengths traces for segments T3–A7, each trace normalized to its overall min. and max. values, traces vertically offset for visibility.

B: Same as A, for different segments of jGC7f larva 18.

C: Example segment length and fluorescence traces for 30 seconds of recording, Gerry larva 19. Top three subpanels: length trace and contraction onsets for the indicated segment in black and magenta, respectively; fluorescence trace and recruitment onsets in green. Both length and fluorescence traces normalized to their overall min. and max. values; length traces inverted for ease of comparison. Bottom subpanel: fluorescence traces for segments T3–A7, each trace normalized to its overall min. and max. values, traces vertically offset for visibility. The example cycle missing fluorescence values from segment A7 was omitted from analysis of recruitment.

All panels: Time windows shown are identical across all four subplots of each panel. Larva numbering consistent with Figures S1, S3, S4, S9.

**Supplemental Figure 3: Phase delay between segmental recruitment and contraction does not depend on larval size.**

For each Gerry larva (n=17), each segment's phase delays between contraction onset and recruitment onset are plotted for each locomotor cycle. Black dots, means. Plots are sorted (left to right, top to bottom) by increasing median segment length as shown in Figure S1; larva numbering consistent with Figures S1, S2, S4, S9.

**Supplemental Figure 4: Individual larval intersegmental coupling of contraction amplitudes, rates, and durations.**

A: For each larva in the full dataset (n=35), mean pairwise intersegmental correlations (Pearson's r) for contraction amplitudes. Correlation values along the diagonal = +1 by construction. Axes (segment ordering) and color scaling identical for all larvae. Pink outline surrounds Gerry larvae.

B: Same as A, for contraction rates.

C: Same as A, for contraction durations.

D-F: Analogous to A-C, for speed-corrected contraction metrics.

**Supplemental Figure 5: Averaged intersegmental kinematic coupling of contraction amplitudes, rates, and durations, before accounting for cycle speed.**

A: Mean pairwise intersegmental correlations for the indicated feature of segmental contraction, without speed correction. Correlation values averaged across n=35 larvae.

B: For each pair of segments, fraction of larvae having significantly (p<0.05) higher-than-expected pairwise correlations, without speed correction.

Color scaling identical across subplots of A and across subplots of B.

**Supplemental Figure 6: Correlations among segments' recruitment features are stronger than among contraction features.**

A: Pairwise intersegmental correlation strength of the indicated recruitment feature, averaged over all segment pairs at each intersegmental distance. Asterisks: correlations significantly different from zero after Fisher's z-transformation, two-tailed T-test, alpha=0.05 with Holm–Bonferroni correction. n=17 larvae.

B: Analogous to A, for speed-corrected recruitment features. Black asterisks: correlations significantly different from zero. Blue asterisks: correlations significantly different from original pre-correction values (cf. panel A).

C: Mean pairwise intersegmental correlations for the indicated feature of segmental recruitment, following speed correction (averaged over n=17 larvae). All values along the diagonal = +1.

D: For each pair of segments, fraction of larvae with significantly (p<0.05) higher-than-expected pairwise correlations, following speed correction.

**Supplemental Figure 7: A posterior block of segments with strongly-correlated contraction durations frequently emerges under a variety of block modeling assumptions.**

A: For best-performing block models of contraction duration correlations, without speed correction (n=35): Number of high-correlation blocks (adjacent-segment r>0.5).

B: Same models as A: Location of the edges of high-correlation blocks, indicating how often blocks ended between a given pair of segments.

C: Same models as A: Frequency with which models included each pair of segments inside the same high-correlation block.

D-F: Analogous to A-C, for best-performing block models of contraction duration correlations (n=35) when considering all blocks (i.e., not thresholding blocks by high adjacent correlation strengths).

G-I: Analogous to A-C, for best-performing block models of contraction duration correlations (n=35) that used different internal correlation assumptions when modeling blocks: rather than decreasing correlations as a function of intersegmental distance within a block, models assumed a flat (equal) correlation strength for all pairs of segments within a block.

**Supplemental Figure 8: Block structure of contraction rate and amplitude correlations.**

A: For best-performing block models of non-speed-corrected contraction rate correlations (top; n=35) and contraction amplitude correlations (bottom; n=35): Number of high-correlation blocks (adjacent-segment r>0.5).

B: Same models as A: Location of the edges of high-correlation blocks, indicating how often blocks ended between a given pair of segments.

C: Same models as A: Frequency with which best-performing models included each pair of segments inside the same high-correlation block.

D-F: Analogous to A-C, for block models of speed-corrected contraction rate and contraction amplitude correlations (n=35 each).

G-I: Analogous to A-C, for best-performing block models (n=35 each) that used different internal correlation assumptions when modeling blocks: rather than decreasing correlations as a function of intersegmental distance within a block, models assumed a flat (equal) correlation strength for all pairs of segments within a block.

**Supplemental Figure 9: Strength of posterior contraction duration correlations tended to decrease with increasing larval width.**

A: Individual larvae's pairwise intersegmental contraction duration correlations. Plots ordered by increasing larval size. Axes (segment ordering) and color scaling identical for all larvae. Pink outline surrounds Gerry larvae.

B: Analogous to A, for correlations calculated using speed-corrected contraction durations.

C: Scatter plot of individual larvae's mean segment width against its ratio of mean posterior to mean anterior contraction duration correlation. Mean P: average of all correlations between pairs of segments A5 through A8. Mean A: same, for pairs of segments T3-A4. n=35 larvae; points colored per larva.

D: Scatter plots of same data as in (C), separated into jGC7f (left) and Gerry (right) subpopulations.
